# Arabidopsis Nicotianamine Synthases (NAS) comprise a common core-NAS domain fused to a variable auto-inhibitory C-terminus

**DOI:** 10.1101/2022.10.13.512114

**Authors:** Hiroyuki Seebach, Gabriel Radow, Michael Brunek, Frank Schulz, Markus Piotrowski, Ute Krämer

## Abstract

Nicotianamine Synthase (NAS) catalyzes the biosynthesis of nicotianamine (NA) from the 2-aminobutyrate moieties of three *S*-adenosylmethionine molecules. NA has central roles in metal nutrition and metal homeostasis of flowering plants. Despite the availability of crystal structures of archaeal and bacterial NAS-like proteins that carry out simpler aminobutanoyltransferase reactions, the enzymatic function of NAS remains poorly understood. Here we report amino acids essential for the activity of AtNAS1 based on structural modeling and site-directed mutagenesis. An enzyme-coupled continuous activity assay allowed us to compare differing NAS proteins identified through multiple sequence alignments and phylogenetic analyses. In most class Ia and b NAS proteins of dicotyledonous and monocotyledonous angiosperm plants, respectively, the core-NAS domain is fused to a variable C-terminal domain. Compared to fungal and moss NAS (class III) that consist merely of the core-NAS domain, NA biosynthetic activities of the four paralogous Arabidopsis NAS proteins were far lower. Yet their C-terminally trimmed core-NAS variants exhibited strongly elevated activities. Out of 320 amino acids of AtNAS1, twelve, 287-TRGCMFMPCNCS-298, accounted for the auto-inhibitory effect of the C-terminus, with approximately one third contributed by N296 within a CNCS motif that is conserved in Arabidopsis. No detectable NA biosynthesis was mediated by two representatives of groups of plant NAS proteins that naturally lack the C-terminal domain, class Ia *Arabidopsis halleri* NAS5, and *Medicago truncatula* NAS2 of class II which is found in dicots and diverged early during the evolution of flowering plants. Our results suggest that NAS activity is under stringent post-translational control in plants.

## Introduction

Maintaining adequate uptake, distribution and storage of essential metals, for example iron (Fe), zinc (Zn) and copper (Cu), is critical for the survival and fitness of all organisms (1, 2). Thus, metal homeostasis networks operate by orchestrating a variety of transmembrane metal transport, metal chelation and metal trafficking processes. In land plants, the non-proteinogenic amino acid nicotianamine (NA) has a central function as a low-molecular-weight chelator molecule which can bind cations of Fe, Zn, Cu, and other metals. Nicotianamine Synthase (NAS, EC 2.5.1.43; Fig. 1 and SI Fig. S1) enzymes, first identified in angiosperm plants, catalyze the biosynthesis of one molecule NA from three molecules of SAM in a step-wise fashion (3–6). In plants, NA can act in the cytosol, vacuole, xylem and phloem to affect the intra-cellular sequestration or the intra-cellular, inter-cellular and long-distance partitioning of metals, often in a localized and at least partially metal-specific fashion through an interplay with transmembrane transporters of differing substrate specificities. For instance, NA acts in the transport of iron from the phloem outwards in sink organs, for example young leaves or developing seeds (5, 7–11). In graminaceous monocotyledons, NA additionally serves as the precursor for the biosynthesis of phytosiderophores, such as 2’-deoxymugineic acid, which are secreted into the rhizosphere for scavenging Fe(III) prior to the uptake of FeIII-phytosiderophore complexes into root cells in strategy II of plant iron uptake (12).

**Figure 1.**
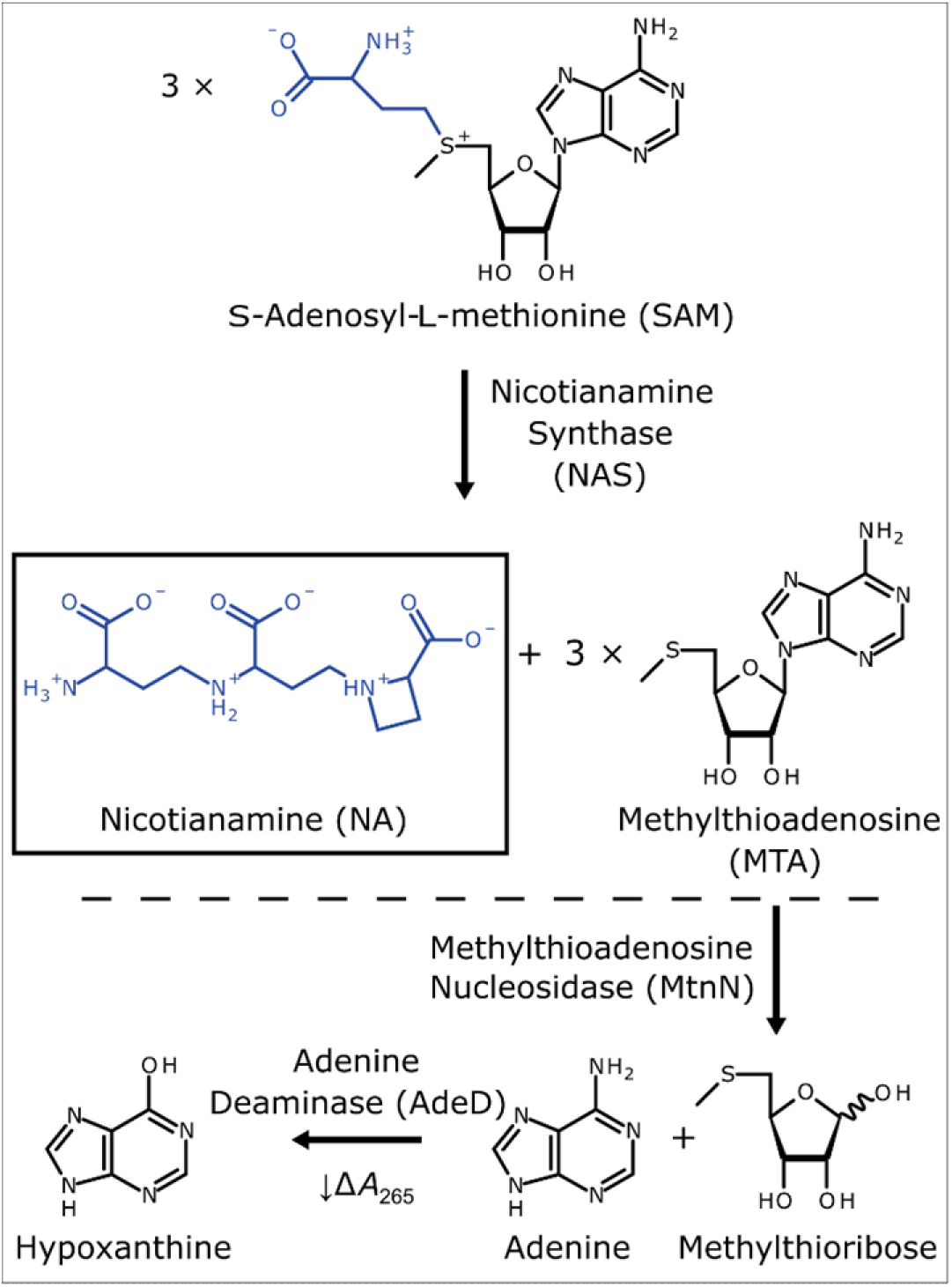
Biochemical function of Nicotianamine Synthase (NAS) and general strategy of the coupled spectrophotometric assay to quantify NAS enzyme activity. Nicotianamine Synthase (NAS) enzymes catalyze the formation of one molecule of nicotianamine from three molecules of SAM (upper part). The byproduct of this reaction, MTA, can be converted to hypoxanthine through two sequential reactions catalyzed by MtnN and AdeD, respectively (lower part). The formation of hypoxanthine can be monitored spectrophotometrically through a decrease in absorbance at 265 nm.

Although initially thought to be unique to seed plants, NAS proteins and NA production were subsequently identified in the moss *Physcomitrium (Physcomitrella) patens* within the division of bryophytes, and in the filamentous fungus *Neurospora crassa* (13–16). Moreover, NAS-like enzymes were described in the bacteria *Staphylococcus aureus*, *Pseudomonas aeruginosa* and *Yersinia pestis*, as well as in the archaeon *Methanothermobacter thermautotrophicus* (17–20). A crystal structure of MtNAS provided seminal insights into its catalytic mechanism (18, 21). Plant and fungal NAS enzymes sequentially use the 2-aminobutyrate moieties of three SAM molecules to form one molecule of NA, releasing three molecules of 5’-methylthioadenosine (MTA) as a byproduct (12) (Fig. 1 and SI Fig. S1). In NAS-like enzymes, various amino acids can apparently serve as the starter molecule for the reaction, namely glutamate, D-histidine or L-histidine (SI Fig. S1). NAS-like enzymes link the α-amino nitrogen of the starter molecule acting as a nucleophile onto the C_4_ atom of a 2-aminobutyrate moiety from SAM (Fig. 1 and SI Fig. S1). Subsequently, there can be an additional cycle of extension, either by using another 2-aminobutyrate moiety from SAM similar to plant and fungal NAS, or alternatively by using pyruvate or α-ketoglutarate (Fig. 1 and SI Fig. S1). As a result, NAS-like enzymes catalyze the formation of NA-like compounds thermoNA (tNA), xNA or yNA (17, 22).

To date, the absence of a sensitive and quantitative assay for the analysis of enzyme activity has hampered biochemical studies of NAS enzymes. Instead, NAS activity was demonstrated upon separation of reaction mixtures either by thin-layer chromatography, for example through autoradiography of [^14^C]-NA formed from [^14^C]-SAM (e.g., (4, 7)), or alternatively by HPLC, with photometric or mass-spectrometric detection of derivatized or underivatized NA (e.g., (13, 23)). NAS enzymes are strongly feedback-inhibited by MTA, which is also a spontaneous breakdown product of the labile substrate SAM (7, 23). Thus, assays were performed in the presence of large amounts of the NAS enzyme and low concentrations of the SAM substrate, therefore requiring sensitive methods to detect the (low amounts of) NA formed.

Here we report that a continuous enzyme-coupled photometric assay enabled us to quantify the catalytic activities of a number of previously characterized and uncharacterized recombinant NAS proteins *in vitro*. Compared to activities between 0.2 and 0.8 nkat (mg protein)^−1^ of fungal and moss NAS, the activity of *A. thaliana* NAS1 (AT5G04950) was less than one tenth and thus substantially lower. To understand the cause of such strikingly different activities of NAS proteins from different organisms, we conducted a phylogenetic analysis of NAS homologs predicted from publicly available nucleotide sequence data. We resolved AtNAS1 to AtNAS4 (class Ia), as well as NcNAS (class III), for example, in their expected relative phylogenetic positions. In addition to the previously characterized dicot (class Ia) and monocot (class Ib) NAS proteins, a new group of NAS proteins (class II) is represented in a subset of dicots. Moreover, a fifth class Ia NAS1/NAS2 paralog, NAS5, is encoded in the genomes of numerous Brassicaceae species including members of the *Arabidopsis* genus, but not *A. thaliana*. Amino acid sequence alignments revealed the presence of an elongated C-terminus in about 90% of class Ia and b NAS proteins, whereas most other NAS and NAS-like proteins, including also NcNAS and PpNAS, consist predominantly of a core-NAS domain. Enzyme activities of AtNAS mutant variants carrying C-terminal deletions suggested an auto-inhibitory role of the elongated C-termini of these class Ia NAS proteins. Within AtNAS1, which is predicted to comprise 320 amino acids in total, we attributed the auto-inhibitory effect to a segment of 12 amino acids at positions 287 to 298 in the elongated C-terminus, and the single replacement of N296 by D resulted in activation to 30% of the maximal activity of the C-terminally truncated AtNAS1 protein.

## Results

### Establishment of a continuous enzyme-coupled photometric NAS assay

Previously published studies provided qualitative evidence for the enzymatic activities of NAS proteins (13, 24). For quantitative comparisons among NAS enzyme activities, we developed an enzyme-coupled photometric NAS activity assay, based on a published method for the quantification of SAM-dependent methyltransferase activity (25). SAM-dependent methyltransferases catalyze transmethylation reactions using SAM as the donor of a methyl group and release SAH as a byproduct. Their enzyme activities were quantified in a coupled enzyme assay, in which the byproduct SAH is converted to hypoxanthine by the sequential action of *S*-Adenosylhomocysteine Nucleosidase (SAHN/MtnN), and Adenine Deaminase (AdeD). It was thus possible to monitor the activity of SAM-dependent methyltransferases photometrically by following the decrease in absorbance at the wavelength of 265 nm, which results from the deamination of adenine to hypoxanthine. Since many bacterial SAHN also accept MTA, the byproduct of NAS, as a substrate, we hypothesized that these coupled reactions could also be employed in a NAS enzyme activity assay (Fig. 1). Thus, we cloned the coding sequences of *mtnN* (encoding 5’-methylthioadenosine/S-adenosylhomocysteine nucleosidase) and *adeD* from the *Escherichia coli* laboratory strain XL-1 blue, overexpressed them in the same strain and purified the enzymes as recombinant His_6_-tagged fusion proteins. Both recombinantly produced enzymes, MtnN and AdeD, were active in NAS reaction buffer when assayed individually with their respective substrates, as shown by thin-layer chromatography (TLC) (SI Fig. S2A and B). Using both enzymes together, the continuous photometric monitoring of the two-step conversion of MTA to hypoxanthine (decreasing) was possible (SI Fig. S2C).

Next, we tested if this two-enzyme system can be employed for the quantification of NAS activity using *A. thaliana* NAS1 (AtNAS1). First, AtNAS1 was co-incubated *in vitro* with MtnN only, and the assay solutions were then analyzed by TLC for the formation of adenine and nicotianamine (Fig. 2A). Importantly, NA and adenine were clearly detectable when active MtnN was added, but not in the absence of MtnN. This is, to our knowledge, the first time that in a small-scale *in vitro* NAS assay NA formation could be demonstrated simply by ninhydrin staining after TLC, without using radioactive labelled SAM or tedious product purification and concentration steps. In addition, this result confirms the strong inhibitory effect of the byproduct MTA on NAS enzyme activity, which was described earlier (7, 23). Finally, newly formed adenine is a sensitive indicator for NAS activity in this coupled assay (but note that small amounts of adenine seem to be present in the samples as degradation product or contamination of SAM, as indicated by the lanes 1, 4 and 5, Figure 2A).

**Figure 2.**
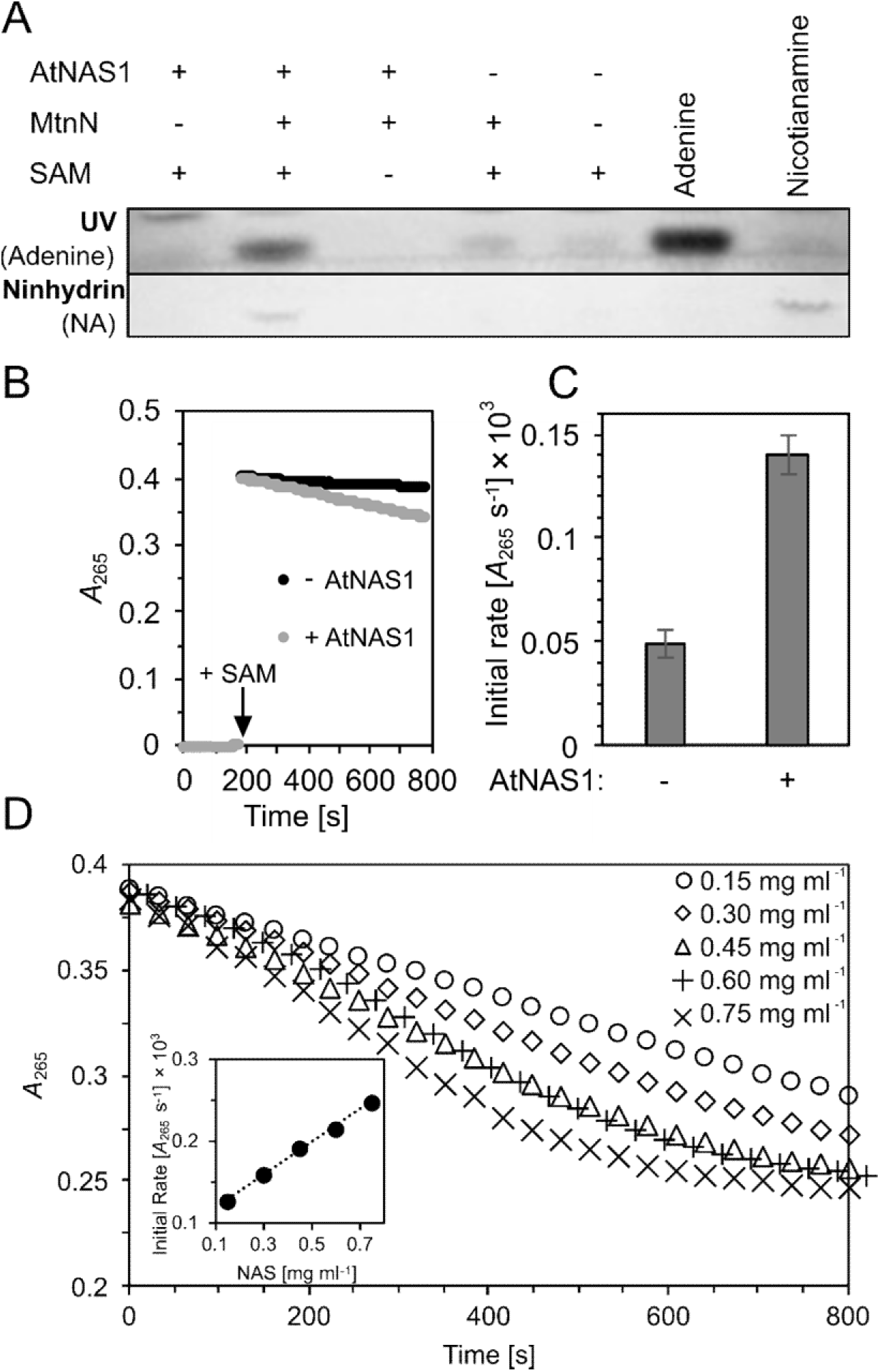
Qualitative and quantitative detection of NAS activity. (A) Images documenting the influence of MtnN on AtNAS1 activity as detected employing the conventional assay. Active (+) or heat-denatured (-) AtNAS1 (0.26 mg ml^−1^) was incubated with active (+) or heat-denatured (-) MtnN (0.1 mg ml^−1^) in the presence (+) or absence (-) of the substrate SAM (5 mM) for 30 min. Five-μl aliquots were separated by thin-layer chromatography (TLC), and the reaction products were visualized in UV light (upper panel) and by ninhydrin staining (bottom panel). Adenine, 14 nmol standard; nicotianamine, 7.5 nmol standard). (B) Coupled spectrophotometric NAS activity assay. MtnN and AdeD were co-incubated with (0.15 mg ml^−1^, light grey) or without AtNAS1 (black) for 3 minutes before the reaction was started by the addition of SAM. (C) Bargraph showing initial rate of change in light absorbance when using 0.15 mg ml^−1^ AtNAS1 (see B) as quantified with the online tool ICEKAT. (D) Coupled spectrophotometric assay conducted with a range of protein concentrations (see legend in diagram) of a truncated AtNAS1 ∆C42 variant lacking the 42 C-terminal amino acids (see Fig. 6 below). The inset demonstrates a linear relationship between protein concentration and the initial rate of change in light absorbance (*y* = 0.1987*x* + 0.01007, *r*^2^ = 0.9985).

This experimental setup allowed us to qualitatively examine the effects of mutations in NAS proteins, which we introduced in order to identify amino acids that are required for the activity of AtNAS1 (Fig. 1 and SI Fig. S1). We thus substituted amino acids corresponding to those suggested to be critical for catalysis in MtNAS (18), i.e. E77 and Y106 of AtNAS1, and amino acids that are conserved among plant NAS proteins but differ between plant NAS and microbial NAS-like proteins, i.e. C69, E73 or the 206-VGMD-209 motif of AtNAS1 (SI Fig. S3). The failure to produce adenine and NA in the assays of the corresponding mutant AtNAS1 proteins E77Q and Y106F suggested that E77 and Y106 are essential for the catalysis of NA formation by AtNAS1. This observation was in agreement with the proposed role of the corresponding amino acids in the formation of tNA by MtNAS, based on crystal structures (18). In addition, mutations introduced at two sites that are conserved only among plant NAS proteins and predicted to be positioned near the reaction chamber, i.e. the replacement of E73 by Q, or the deletion of the 206-VGMD-209 motif, also rendered AtNAS1 inactive, whereas AtNAS1 activity was insensitive to the C69A substitution (SI Fig. S3C).

Next, we additionally included AdeD in our assay to test whether this allows to follow the NAS reaction photometrically. In the presence of AtNAS1, MtnN and AdeD, there was a measurable decrease in light absorbance at an initial rate that was about three times higher than the background in the absence of AtNAS1 and that remained constant over several minutes (Fig. 2B, C). By comparison, the minor decrease in light absorbance of a reaction mixture containing the enzymes AdeD and MtnN alone may result from the presence of low levels of contaminating adenine in the SAM solution (see Fig. 2B). Importantly, there was a linear correlation between the concentration of the NAS protein in the assay and the initial rate quantified according to the change in *A*_265_ over time, indicating against a limitation by insufficient amounts of MtnN and AdeD present in our assay (Fig. 2D).

### Proof of concept for a one-pot *in vitro* synthesis of nicotianamine from ATP and methionine

Previously described systems for the biosynthetic production of NA used either crude extracts from plants or recombinant NAS proteins from plants or *N. crassa* in yeast cells (23, 26–28). We asked whether we could take advantage of the increased activity of AtNAS1 in the presence of MtnN (Fig. 2A) for the *in vitro* production of NA. Combining MetK, a bacterial SAM synthetase which synthesizes SAM from ATP and l-methionine, with both AtNAS1 and MtnN should allow the biosynthesis of NA from these precursors. Initially, we cloned the *E. coli metK* gene, overexpressed it and purified the enzyme as His_6_-tagged fusion protein for the production of fresh SAM in order to replace the (presumably partly degraded) commercially available SAM in our NAS assays. Subsequently, we combined MetK with MtnN and AtNAS1 for the one-pot synthesis of NA from ATP and methionine (SI Fig. S4). We approximated that at least 60 nmol (18 μg) NA were produced within 4 h in a volume of 120 μl in the presence of 60 μg MetK, 12 μg MtnN and 22 μg AtNAS1. In the past, 4.5 nmol (1.3 μg) NA was produced by 350 μg NASHOR1 (a NAS from *Hordeum vulgare*) using SAM as a substrate, 60 to 750 μg [^15^N_3_]-NA was obtained per 120-ml culture of recombinant *Schizosaccharomyces pombe* cells producing NcNAS, and an engineered strain of *Saccharomyces cerevisiae* producing AtNAS2 yielded 766 μg NA g^−1^ wet biomass (23, 26, 28).

### Phylogenetic analysis and sequence comparison

NAS proteins are encoded in the genomes of some archaea, bacteria, fungi, mosses, and all land plants. To understand how the amino acid sequences of different NAS proteins are related among one another, we conducted a phylogenetic analysis. We constructed a phylogenetic tree by Bayesian inference based on the shared core region of 186 NAS and NAS-like proteins, which we defined as the part of the multiple sequence alignment corresponding to the segment from the 1^st^ and 275^th^ amino acid of AtNAS1 in order to exclude any N- or C-terminal extensions present in only a subset of proteins.

NAS proteins from dicotyledonous plants grouped in two distinct monophyletic clades with high statistical support. One of these clades comprised the well-known NAS proteins (termed here class Ia), for example those of *A. thaliana* and tomato, and it was the sister clade of the NAS proteins from monocotyledonous plants (class Ib), of which barley the HvNAS proteins were characterized first, for example (5, 7, 29). The second, distinct clade of NAS proteins from dicotyledonous plants (class II), comprised annotated NAS proteins from the Ranunculaceae family of basal eudicots, the Apiaceae family in the Asterid clade, as well as the Rutaceae, Rosaceae, Fabaceae, Salicaceae, Euphorbiaceae and Malvaceae from the Rosid clade. Clade credibility values supported only weakly that class II NAS proteins diverged from class I NAS proteins before the origin of extant gymnosperm NAS proteins (class Ic) and NAS proteins of the basal angiosperm *Amborella trichopoda* (class I/Ic). With stronger support, class II NAS proteins diverged from class I NAS proteins before monocot and dicot class I NAS proteins diverged from one another. NAS from fungi and mosses (class III), archaea (class IV) and bacteria (class V) diverged earlier from all Spermatophyte NAS proteins, consistent with a published phylogeny of a small set of proteins (15) (Fig. 3).

**Figure 3.**
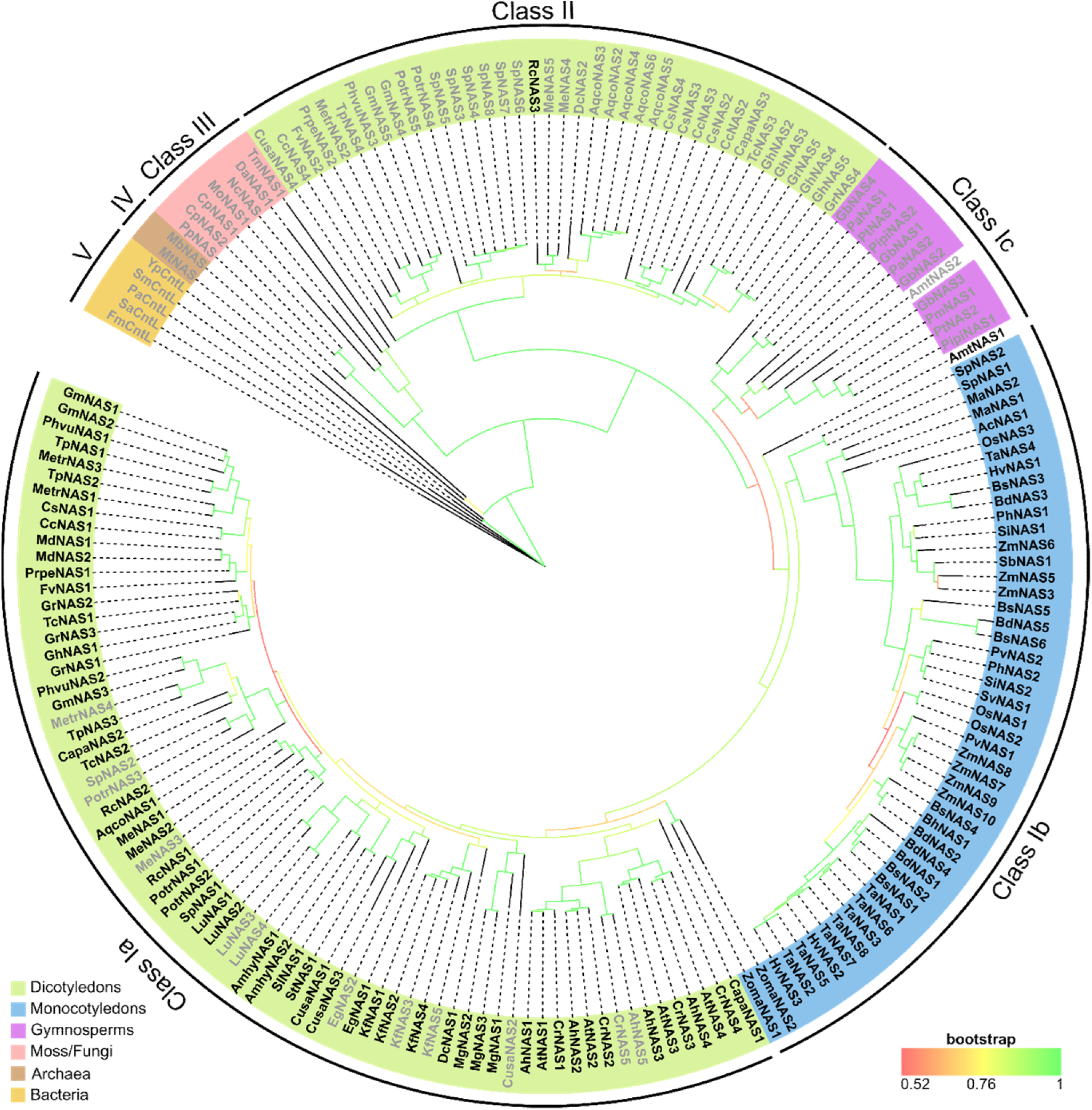
Phylogenetic analysis of plant NAS proteins. Phylogenetic tree of NAS proteins from seed plants, mosses, and fungi, as well as NAS-like proteins from bacteria and archaea (SI Table S1) obtained by Bayesian inference. Short NAS (generally about 280 amino acids in length) end after the core-NAS domain (protein names given in light grey fonts). Long NAS (generally about 320 amino acids in length) comprise additional amino acids at the C-terminus in almost all cases (protein names given in black fonts). Colored backgrounds of short protein names reflect taxonomic groups. Colors of lines at branch positions reflect clade credibility values between 0.52 (red) and 1 (green).

Our phylogenetic analysis further suggested that the amino acid sequences of the NAS proteins from the mosses *P. patens* and *Ceratodon purpureus* are more closely related to fungal than to plant NAS proteins. This was further supported by the fact that both fungal and moss *NAS* genes contain an intron at a conserved position that also conserves the phase of the intron in relation to the codons of the coding sequence (SI Fig. S5), whereas all *NAS* genes from seed plants are intron-free. The genomes of the mosses *Marchantia polymorpha*, *Sphagnum fallax* and *S. magellanicum* do not contain any *NAS* genes. These observations support that the *NAS* gene of *P. patens* and *C. purpureus* is of fungal origin and probably arose through a horizontal gene transfer.

Multiple sequence alignments revealed that all class Ib and many of the class Ia NAS proteins have an extended C-terminus of approximately 40 aa (Fig. 4, termed here long NAS, length of about 320 aa), in contrast to almost all other proteins in class Ic and classes II to V (termed here short NAS, length of about 280 aa)(Fig. 3). The basal angiosperm *A. trichopoda* possesses both a long and a short NAS protein, with the latter positioned at the base of class Ia and Ib. Our analysis suggested that there were several independent secondary losses of the extended C-terminus in class Ia, as exemplified by the NAS isoforms MetrNAS4 and AhNAS5 (Figs. 3 and 4).

**Figure 4.**
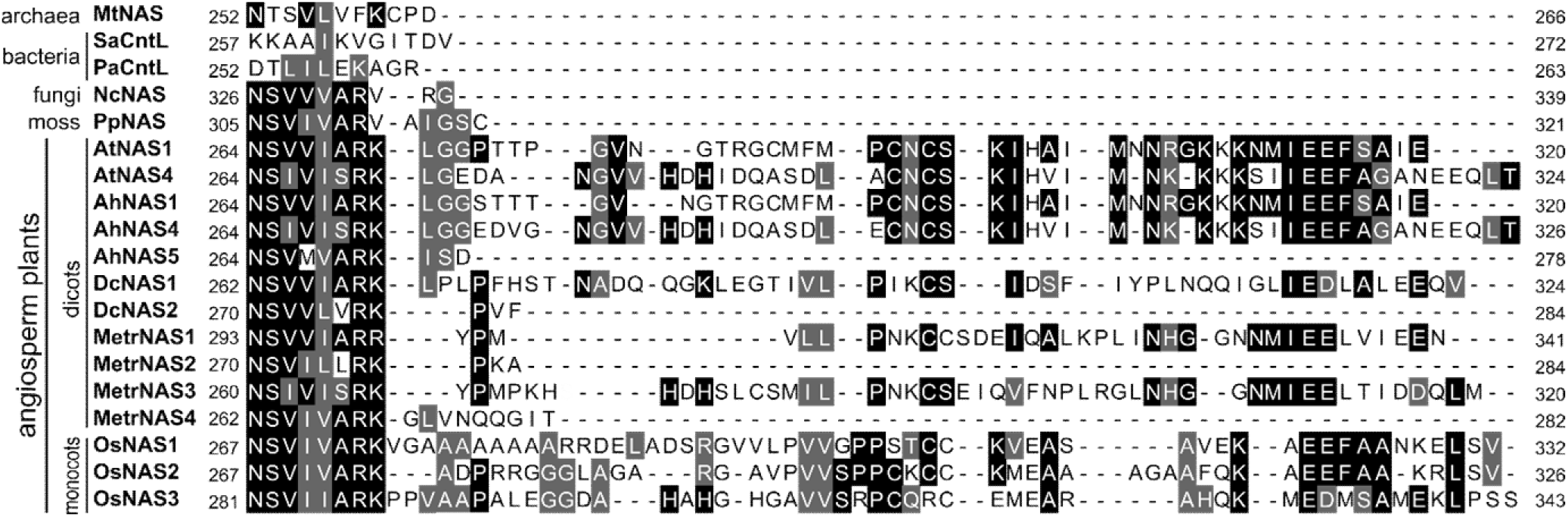
Alignment of amino acid sequences of C-terminal regions of NAS and NAS-like proteins from archaea, bacteria, fungi, and plants. Amino acids are shown on a black/grey background whenever ≥ 50% of them are identical/similar, considering only sequences ungapped at the respective position. Similarity groups were based on the default classification of MultipleAlignShow (ILV, FWY, KRH, DE, GAS, P, C, TNQM)(50). The alignment was carried out in MegaXI using ClustalW with standard settings. Note that among the angiosperm plant NAS, AhNAS5 (class Ia), DcNAS2 (class II), MetrNAS2 (class II) and MetrNAS4 (class Ia) are short NAS, whereas the remainder are long NAS. Mt: *Methanothermobacter thermautotrophicus*, Sa*: Staphylococcus aureus*, Pa: *Pseudomonas aeruginosa*, Nc: Neurospora *crassa*, Pp: *Physcomitrium patens*, At: *Arabidopsis thaliana*, Ah: *Arabidopsis halleri*, Dc: *Daucus carota*, Metr: *Medicago truncatula*, Os: *Oryza sativa* (SI Table S1).

### Auto-inhibitory effect of the elongated C-terminus in plant NAS enzymes

We quantified the activities of a set of short and long NAS proteins using the coupled photometric assay *in vitro*. Although we had initially used AtNAS1, a long NAS, to establish the enzyme activity assay, its activity was only very low (0.02 ± 0.01 nkat (mg protein) ^−1^ in two independent preparations). By contrast, the activities of NcNAS (0.81 ± 0.01 and 0.51 ± 0.05 nkat mg^−1^ protein in two independent preparations) were the highest, followed by PpNAS (0.37 ± 0.03 and 0.21 ± 0.01 nkat mg^−1^ protein), both of them being short NAS proteins (Fig. 5). To our knowledge, the activity of the NAS enzyme from the moss *P. patens* had not been demonstrated earlier. The biosynthesis of NA was confirmed through mass spectrometry (SI Fig. S6). These data suggested the possibility of an influence of the C-terminal domain on the activity of NAS proteins.

**Figure 5.**
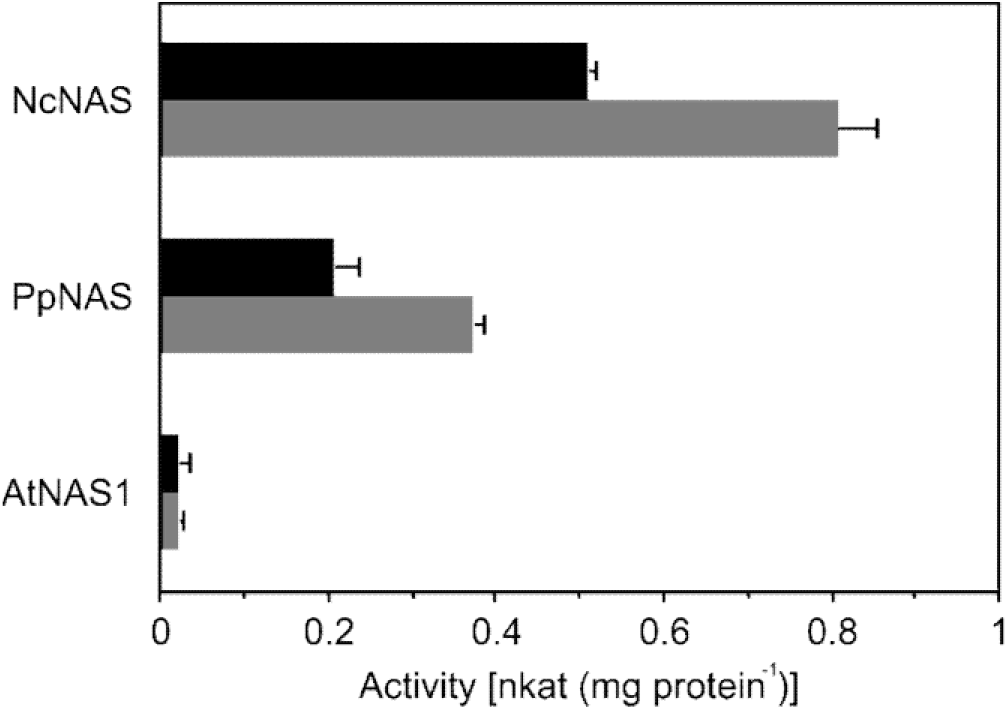
Comparison of enzyme activities of NAS from different species. Enzyme activities were quantified in the coupled spectrophotometric assay (see Fig. 2B, D). Bars show mean ± SD (*n* = 3 technical replicates) from two independent protein purifications (black and grey, respectively). At: *Arabidopsis thaliana*, Nc: *Neurospora crassa*, Pp: *Physcomitrium patens*.

As a representative of a new group of NAS enzymes in the Brassicaceae and an example of a short NAS of secondary origin in class Ia, we tested AhNAS5 activity *in vitro*. However, its activity, as well as any formed NA, were below our detection limits (SI Fig. S6). Given the conserved sequence changes and the wide distribution of NAS5 homologs in the Brassicaceae, a neo-functionalization for a different, yet unidentified, function is possible (SI Fig. S7). It should be mentioned that the plant model organism *A. thaliana* has lost most of its *NAS5* gene, but a remaining segment of it encoding a partial protein homologous to the 57 C-terminal amino acids of AhNAS5 is still present in the genome (AGI code AT4G26483). The encoded protein is predicted to have a length of 84 aa and is unlikely to have any NAS or NAS-related activity (SI Fig. S7C). Furthermore, we tested the *in vitro* activity of MetrNAS2, a short NAS of class II. Similar to AhNAS5, both MetrNAS2 activity as quantified in our photometric assay and the levels of formed NA as analyzed by LC-MS remained below our detection limits (SI Fig. S6).

In order to address a possible influence of the elongated C-terminus on the enzyme activity of plant long NAS proteins, we generated C-terminally truncated variants lacking this domain and containing only the core-NAS domain for each of the four NAS homologs of *A. thaliana* and quantified their enzyme activities (Fig. 6A, B). As for AtNAS1, the enzyme activities of the full-length AtNAS2 to AtNAS4 were low or even around the limit of detection. C-terminal truncation caused a strong, 30- and 20-fold increase in the activities of AtNAS1 and AtNAS2 *in vitro*, respectively. The truncated variants of AtNAS3 and AtNAS4 also showed strongly increased activities that remained below 25% of those of C-terminally truncated AtNAS1 and AtNAS2 (Fig. 6B).

**Figure 6.**
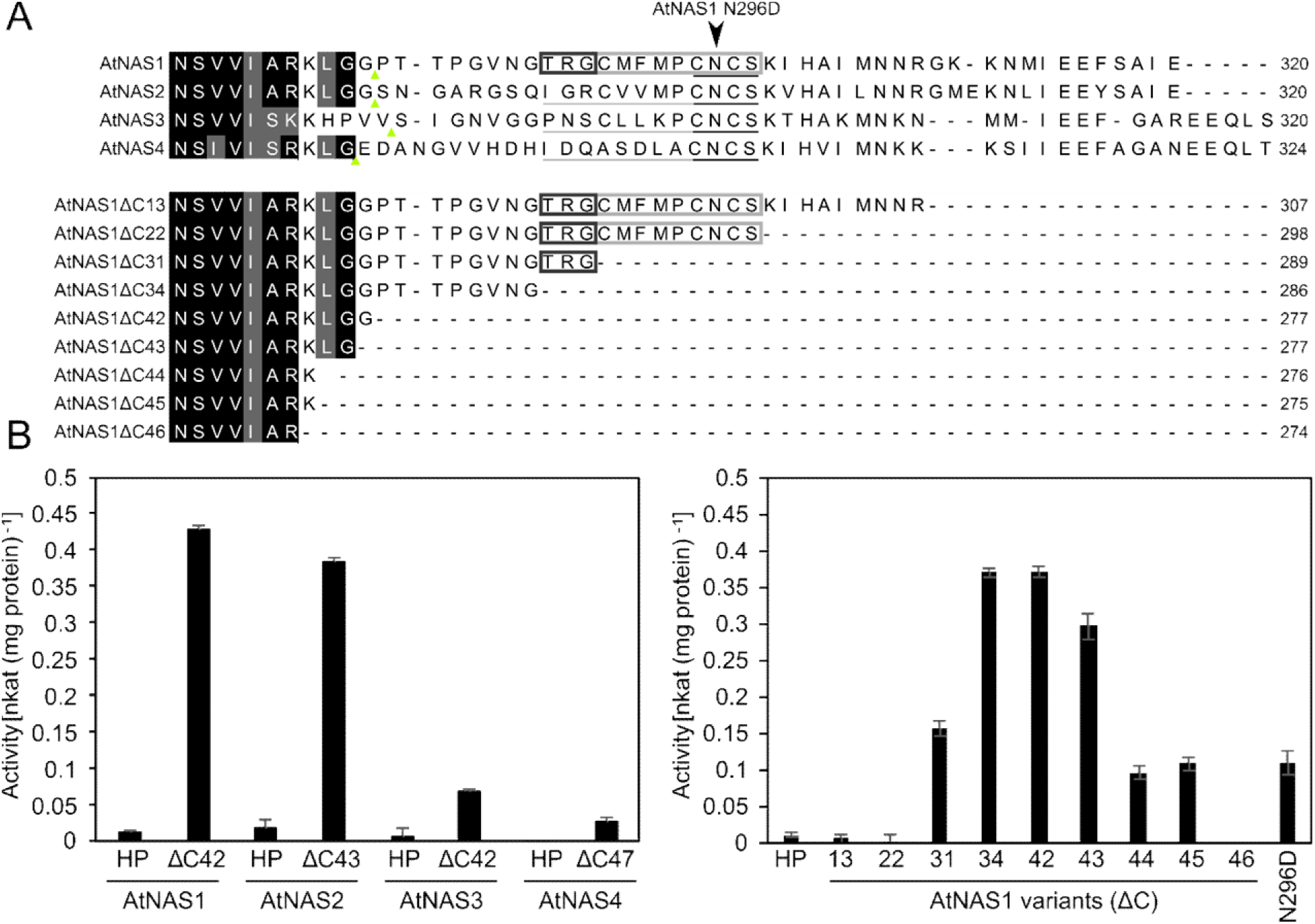
Inhibitory effect of the variable C-terminal domain on the activity of Arabidopsis NAS proteins. (A) Alignment of AtNAS1, −2, −3 and −4 (upper part) and stepwise C-terminally truncated AtNAS1 variants (bottom part). Numbers following ∆C indicate the number of amino acids deleted from the C-terminus of AtNAS1. Amino acids fully conserved/similar by comparison to short NAS PpNAS and NcNAS (or one of them) are highlighted in black/grey (see Fig. 4). (B) Activity of full-length and C-terminally truncated variants for the four paralogous NAS proteins of *A. thaliana*. The C-terminal end of each truncated variant is marked by a small grey triangle in (A, upper part). (C) Effect of progressive C-terminal deletions on AtNAS1 activity. Bars show mean ± SD (*n* = 3 technical replicates). HP: Holoprotein.

Next, we analyzed the effects of progressive C-terminal deletions on the activity of AtNAS1 (Fig. 6A, lower part, and 6 C). Removal of the C-terminal 13 or 22 amino acids of AtNAS1 did not result in any increase in its enzyme activity. Deletion of 31 amino acids resulted in 42% of the maximal activity that was observed upon deletion of either 34 or 42 amino acids from the C-terminus. Compared to the maximal activity, AtNAS1 activity decreased again when it was C-terminally truncated by more than 42 amino acids. C-terminal truncation by 46 residues removed a lysine residue that is highly conserved among plant NAS proteins, and this eliminated enzyme activity. Thus, we attributed the repression of AtNAS1 activity to 12 amino acids, residues 287 to 298 of the C-terminus. The corresponding amino acid sequence, TRGCMFMPCNCS, is partly conserved between AtNAS1 and AtNAS2-4, with full conservation of the final 4 amino acids, CNCS. Of the inhibitory effect, about 58% appeared to reside in the first three, non-conserved amino acids 287-TRG-289 of the 12-amino-acid auto-inhibitory region.

We next tested the effects of exchanging single amino acids between positions 290 and 298. Out of the variants C290A, F292L, P294A, C295A, N296D, C297A and S298A of full-length AtNAS1, only N296D exhibited an activation to 30% of the maximal activity of C-terminally truncated AtNAS1 (AtNAS ∆42) (Fig. 6C).

## Discussion

A major problem in the biochemical characterization of NAS enzymes had been the low enzyme activity so far, which was probably caused by a strong inhibitory effect of the byproduct MTA on the reaction catalyzed by the NAS enzyme (7, 23). To alleviate this, researchers had used only low concentrations of the substrate SAM (20 μM) in NAS activity assays, which, in turn, required sensitive methods to detect the small amounts of NA formed. MTA may already be present as contaminating or breakdown product in commercially available SAM preparations, further interfering with the sensitivity of conventional assays for NAS activity. Here we report that the inclusion of *E. coli* Methylthioadenosine Nucleosidase (MtnN) in the NAS assay, which irreversibly hydrolyzes MTA to adenine and methylthioribose, substantially increased the amount of NA formed by AtNAS1 in an endpoint assay (Fig. 2A). We attribute this effect to the degradation of newly formed MTA by MtnN, which counteracts a gradual accumulation of MTA over time in the reaction mix and thus prevents the inhibition of NAS by MTA. Consequently, the combination of NAS with MtnN allows a simple, semi-quantitative NAS assay, in which many samples can be analyzed in parallel for the formation of NA and adenine after separation of the reaction products by TLC (SI Fig. 3C). By further adding Adenine Deaminase (AdeD), which converts adenine to hypoxanthine, the NAS reaction can be continuously monitored by spectrophotometry (Fig. 2B-D). It should be noted that the activities of poorly active NAS enzymes, for example the full-length AtNAS1, remain close to the detection limit of this spectrophotometric assay. In the future, it might be possible to further increase the detection limit by including a xanthine oxidase, which acts on hypoxanthine to release hydrogen peroxide for monitoring by colorimetric or fluorometric detection methods using additional reagents. This principle is used in commercial kits for quantifying the activities of SAM-dependent methyltransferases (30).

The strong inhibitory effect of MTA on NAS enzymes *in vitro* raises the question of whether this is of physiological relevance *in planta*. MTA is not only a byproduct of NA biosynthesis, but also of ethylene and polyamine biosynthesis. Plants possess MtnN orthologues which are part in the methionine salvage pathway. Two *MTN* genes reside in the genome of *A. thaliana*, but co-expression of *MTN* and *NAS* genes has not been reported, which may suggest that the basal MtnN activity is sufficient to prevent the accumulation of an excess of MTA in the cytosol. A coordinated upregulation of *NAS* and *MTN* genes under iron deficiency was described in rice and wheat (31–33). NA biosynthetic rates might be higher in these plants than in *A. thaliana* because – being strategy II plants – they use NA also as a precursor for the biosynthesis of phytosiderophores.

A sequence alignment and a phylogenetic analysis including 186 NAS amino acid sequences from a variety of species revealed the existence of two different types of NAS in land plants. Long NAS proteins, found almost exclusively in class Ia and b, differ from short NAS by an additional variable domain at the C-terminus which consists of approximately 40 amino acids. Monocot plants possess only long NAS (class Ib) whereas both long and short NAS are found in dicot plants (class Ia and II). Interestingly, class II that exclusively contains short NAS proteins from dicots, appears to have arisen before the divergence of the monocot and dicot lineages. Note that long NAS of class Ia and b exhibit some shared C-terminal sequence features (see Fig. 5), which were absent in the single class II long NAS protein, *Ricinus communis* NAS3. Indeed, the global multiple sequence alignment clarified that this protein carries an N-terminal instead of a C-terminal extension (SI Data S1). Our phylogenetic analysis clearly indicated that the earliest NAS were short NAS proteins and that NAS proteins containing an extended C-terminus likely arose early during angiosperm evolution. This was supported by the fact that only short NAS proteins are found in archaea, bacteria, fungi or moss. Moreover, our phylogenetic analysis suggests that class II NAS were secondarily lost at least in the monocots and several clades of the dicots. Alternatively, class II may have arisen at an early stage during the evolution of the dicot lineage through horizontal gene transfer from an unknown eukaryotic organism. This remains to be examined in more detail.

We observed a striking difference in activity between a long NAS, namely full-length AtNAS1, and its C-terminally truncated variant comprising only the core-NAS domain, AtNAS1 ∆C42, which exhibited a much higher *in vitro* enzyme activity (Fig. 6). We obtained similar results for AtNAS2, AtNAS3 and AtNAS4. In accordance with this, enzyme activities of the short NAS proteins PpNAS and NcNAS were also much higher than that of AtNAS1 (Fig. 5). The NAS substrate SAM is an important methyl donor in a variety of cellular biochemical reactions, for example the methylation of DNA and RNA, and SAM is also a substrate for the biosynthesis pathway of polyamines and of the phytohormone ethylene (34, 35). Consequently, the consumption of SAM by NAS might require tight regulation. Therefore, it is possible that the C-terminus of long NAS in class Ia and Ib also has an auto-inhibitory role *in planta*, and that post-translational mechanisms are required to activate these NAS proteins. The C-terminus contains a noticeably large number of amino acids that can act in the coordination of bound metal ions, such as cysteine, histidine, glutamate or methionine. It is thus possible that these NAS proteins are activated upon the binding of metal cations to, or upon the loss of bound metal cations from, the C-terminal extensions. *In planta*, NAS activation could operate through metal- or protein-binding-dependent or -independent alterations in three-dimensional structure, post-translational modification, or proteolytic processing (36, 37). The *in planta* biosynthesis of NAS protein in an enzymatically inactive form and its controlled post-translational activation would allow a tight control of NAS activity, and thus of SAM consumption and MTA production by NAS. Future work will address these possibilities.

In the bacterial NAS-like protein *Staphylococcus aureus* CntL that comprises merely a core-NAS domain, Luo *et al.* (38) identified a linker region between its N- and C-terminal domain which is capable of conformational alternation between the open and closed states of SaCntL. The authors suggest that this linker region could have pivotal roles in allowing substrate entry, substrate recognition, and catalysis. The linker region corresponds to the amino acids between H96 and P125 of AtNAS1 (SI Fig. S8A). A comparison of a crystal structure of SaCntL with the structure of AtNAS1 as predicted by AlphaFold 2 suggests that the putative linker region is within reach of the C-terminal extension of AtNAS1 (SI Fig. S8B; (39)). We observed an increase in enzyme activity of the N296D variant of AtNAS1 (Fig. 6C). Since asparagine can act differently from aspartate in the formation of hydrogen bonds, it is possible, for example, that N296 binds to the linker region of AtNAS1 *via* hydrogen bonding. As a consequence, the movement of the linker region might be impaired, thus interfering with the catalytic cycle of AtNAS1. Alternatively, hydrogen bonding of N296 might occur to a different region of the NAS protein surface. Further research is required to identify the mechanism of N296-dependent auto-inhibition, and of how additional C-terminal amino acids contribute to auto-inhibition of AtNAS1.

As a representative of Brassicaceae NAS5 proteins, AhNAS5 was inactive in the biosynthesis of NA from SAM despite the absence of an elongated C-terminus. Based on an amino acid sequence comparison of AhNAS5 with several NAS and NAS-like proteins, there are replacements of conserved amino acids, for example AtNAS1 C69 to R and E73 to V in AhNAS5 (SI Fig. S3A and S7). Here we observed that E73Q, but not C69A, eliminates AtNAS1 activity (SI Fig. S3C). A protein model of AhNAS5 superimposed onto the known protein structure of MtNAS positions these amino acids near the reaction chamber indicating the possibility that these mutations render AhNAS5 inactive (SI Fig. S3B). Further studies on AhNAS5 could test the activity of AhNAS5 R69C and V73E mutant variants or even address a possible neo-functionalization of AhNAS5 in the biosynthesis of an NA-related molecule.

To our knowledge, the only functionally characterized class II NAS (without a C-terminal extension) is *Medicago truncatula* NAS2, which is required for symbiotic nitrogen fixation (40). A *MetrNAS2* loss-of-function *Tnt1* insertion mutant had no growth defect under non-symbiotic conditions. Importantly, overall NA content was not significantly different from the wild type in either shoots, roots or nodules of the mutant. It is thus possible that MetrNAS2 catalyzes the biosynthesis of an NA-related molecule *in vivo* or possesses a different biological role.

Site-directed mutagenesis of AtNAS1 indicated that amino acids E77 and Y106, which are conserved by comparison to MtNAS, are necessary for AtNAS1 activity. Similarly, we observed that E73 and 206-VGMD-209, which are conserved among plant NAS proteins and predicted to localize near the reaction chamber, are required for AtNAS1 activity. Earlier work had described a di-leucine motif at positions 112/113 of OsNAS2 as essential for *in vitro* activity (41) (see SI Fig. S3A). These amino acids correspond to positions 116/117 of AtNAS1, which were not examined in this study. Mutation of 107-YVNL-110 of OsNAS2 to AVDL was reported not to affect the *in vitro* activity, but to interfere with the *in planta* activity, of OsNAS2 (41) (see SI Fig. S3). The mutated motif includes the position corresponding to AtNAS1 Y106, which is critical for *in vitro* activity according to the results presented here. Although these results are overall similar, the differences between our findings on AtNAS1 and earlier findings on OsNAS2 will require further study.

In summary, we report here a continuous assay suitable for quantifying NAS enzyme activities. We identify an extended C-terminus present in class Ia dicot and class Ib monocot NAS proteins, but not in other classes of NAS proteins. We show that this extended C-terminus has an auto-inhibitory function in all class Ia NAS proteins of *A. thaliana in vitro*. By contrast, activities of fungal and moss NAS proteins that lack the C-terminal extension are much higher. We show that class II NAS proteins are present in a number of dicots in addition to class Ia NAS proteins. Class II diverged from class I NAS proteins before the divergence of class I into monocot (Ib) and dicot (Ia) NAS proteins, and class II NAS proteins lack a C-terminal extension. Our results pinpoint the possibility of post-translational regulation of NAS activity *in planta* and provide both methods and a classification of NAS proteins that will facilitate their future functional characterization.

## Experimental procedures

### Protein modelling

The predicted protein model of AtNAS1 by AlphaFold 2 was identified by searching for the Uniprot number of AtNAS1 (Q9FF79)(39). The predicted model of AhNAS5 was calculated by SWISS-MODEL (42) with default settings, since no model was available in AlphaFold 2. All protein models were visualized and further modified in PyMOL (Schrödinger LLC, Version 2.1.1).

### Multiple sequence alignments and phylogenetic analyses

The amino acid sequence of Nicotianamine Synthase 1 (NAS1) of *Arabidopsis thaliana* (AT5G04950) was used in a blastp search against all proteins from annotated genomes in Phytozome (version 12.1.5)(43). In total, 164 full-length NAS sequences from 52 plant species were retrieved using an Expect value (*E*) cut-off of 10^−90^. Likewise, four NAS proteins were identified in the Ginkgo Database (https://ginkgo.zju.edu.cn/) (44) and seven NAS proteins in the PLAZA 5.0 database (https://bioinformatics.psb.ugent.be/plaza/) (45). Further amino acid sequences of NAS and NAS-like proteins from bacteria, archaea, and fungi were included based on published data (13, 15, 18, 46–48). While NAS of *Magnaporthe oryzae* (XP_003719353.1) was studied in a published phylogenetic analysis (15), NAS of *Diaporthe ampelina* (KKY38707.1) was newly identified here as described above and included to obtain a more robust tree. A complete list of all NAS and NAS-like proteins used in our analysis can be found in the SI Table S1, with additional information.

The multiple sequence alignment for the phylogenetic tree shown in Fig. 3 was conducted in Mega11 (49) using ClustalW with standard settings. The full alignment consisting of 603 sites (including gaps) was subsequently trimmed at the N- and C-terminal ends to remove highly divergent and gap-rich regions. The final alignment comprised 412 sites, corresponding to the amino acids 1 to 275 of AtNAS1 (SI Data S1), and displayed using Multiple Align Show (50). Phylogenetic analysis was performed using the program MrBayes (version 3.2.1) (51) with a mixed amino acid rate matrix in two independent Markov Chain Monte Carlo analyses for 2 million generations each. The burn-in was set to 25% and a 50% majority rule tree was obtained. The tree was visualized in iTOL (version 5)(52).

NAS5 sequences of *A. thaliana*, *A. halleri*, *A. lyrata*, *Boechera stricta*, *Brassica rapa*, *Capsella rubella*, *C. grandiflora* and *Eutrema salsugineum* were identified in phytozome by blastp using the amino acid sequences of AtNAS1 and AhNAS5 as queries against the genome of each species. The multiple sequence alignment was conducted in ClustalW with standard settings. Phylogenetic analysis was conducted, and the tree visualized, using the program MegaX based on the full alignment. The phylogenetic tree was calculated using the maximum likelihood method and JTT matrix-based model with 500 bootstrap replicates (53).

### Constructs for recombinant protein production

The open reading frames (ORFs) encoding S-Adenosylmethionine Synthase (MetK, EC 2.5.1.6), 5’-Methylthioadenosine Nucleosidase (MtnN, EC 3.2.2.9) and Adenine Deaminase (AdeD, EC 3.5.4.2) were amplified by PCR from genomic DNA (gDNA) of the *Escherichia coli* strain XL1-Blue, omitting the stop codon (SI Tables S2 to S4). Amplification of *metk* and *mtnN* were carried out with DreamTaq Polymerase (Thermo Fisher Scientific, Waltham, US) according to manufacturer’s instructions. Amplification of *adeD* was carried out using Phusion Polymerase (Thermo Fisher Scientific) according to manufacturer’s instructions. For T/A cloning in to pGEM-T Easy vector (SI Table S2), the *adeD* PCR product was additionally gel-purified (NucleoSpin Gel and PCR Clean-Up Kit, Macherey-Nagel, Düren, DE), and 100 ng of purified DNA was subsequently incubated with 0.2 M dATP and 5 U DreamTaq Polymerase (Thermo Fisher Scientific) in DreamTaq buffer (10 μL total volume) according to manufacturer’s instructions. All ORFs were cloned into the pGEM-T Easy vector according to manufacturer’s instructions. Subsequently, sequences were subcloned into pET-21b (+) (SI Table S2) in front of the C-terminal His_6_-tag using the NdeI and XhoI restriction sites, and the construct were verified by Sanger sequencing.

The genomic sequence encoding *Methanothermobacter thermautotrophicus* NAS (MTH675) was amplified by PCR from a pET101 vector (SI Table S2) provided by the lab of Pascal Arnoux (CEA, Saint Paul lez Durance, France), using Phusion Polymerase (Thermo Fisher Scientific) according to manufacturer’s instructions (SI Table S3 and S4). The amplicon was cloned directly into pET-21b (+) as described above using the NdeI and NotI restriction sites.

Three DNA sequences encoding *Neurospora crassa* NAS (XP_958379), *Physcomitrium patens* NAS (XP_024383030) and *Medicago truncatula* NAS2 (MetrNAS2, Medtr2g070310), all codon-optimized for *E. coli* and flanked by the restriction sites NdeI and XhoI, were synthesized by Invitrogen and provided in pMA-RQ vectors (SI Tables S2 and S5). pMA-RQ-NcNAS, pMA-RQ-PpNAS and pMA-RQ-MetrNAS2 were digested with NdeI and XhoI, and inserts subsequently ligated into pET-21b (+) as described for *mtnN* and *adeD* above.

Coding sequences of *NAS1* (AT5G04950) and *NAS4* (AT1G56430) were amplified from cDNA (Omniscript RT Kit, Qiagen) of *A. thaliana* (Col-0) by PCR using DreamTaq Polymerase (Thermo Fisher Scientific), subcloned first into pGEM-T Easy and subsequently into pET-21b (+) using the NdeI and NotI restriction sites as described above (SI Tables S2 to S4). *AtNAS2* (AT5G56080) and *AtNAS3* (AT1G09240) coding sequences were amplified from genomic DNA of *A. thaliana* (Col-0) by PCR using Phusion Polymerase and cloned into pET-21b (+) using the NdeI and NotI restriction sites as described above.

Point mutations were introduced into AtNAS1 according to the QuikChange protocol from Stratagene modified as described in Zheng et al. 2004 (54). The previously generated pET-21b-AtNAS1 construct was used as a template. The amplification of AtNAS1 C69A, AtNAS1 E73Q and AtNAS1 ∆VGMD were carried out using KAPA Hifi-Polymerase (PeqLab Biotechnologie, Erlangen, DE) according to manufacturer’s instructions (SI Table S3 and S4). AtNAS1 E77Q, AtNAS1 Y106F and AtNAS1 N296D were generated using a Phusion Polymerase (Thermo Fisher Scientific) according to manufacturer’s instructions (Tables S3 and S4). The ORFs encoding C-terminally truncated variants of AtNAS1, AtNAS2, AtNAS3 and AtNAS4 were amplified by PCR using Phusion Polymerase (Thermo Fisher Scientific), the same forward primer and different reverse primers according to manufacturer’s instructions (SI Tables S3 and S4). The amplicons for AtNAS1, all C-terminally truncated variants of AtNAS1, AtNAS4 and AtNAS4 ∆C47 were cloned directly into pET-21b (+) using the NdeI and NotI restriction sites as described above. Amplicons encoding AtNAS2 ∆C43 and AtNAS3 ∆C42 and their respective full length ORFs were cloned into pET-21b (+) using the NheI and NotI restriction sites. The ORF of *AhNAS5* was amplified from gDNA of *Arabidopsis halleri* (Lan 3.1) via PCR using Phusion Polymerase (Thermo Fisher Scientific) according to manufacturer’s instructions (SI Tables S3 and S4). The amplicon was subcloned first into pGEM-T Easy and subsequently into its destination vector pET-21b (+) as described above using the NdeI and NotI restriction sites (SI Table S2).

### Production and purification of recombinant proteins

We expressed *metK*, *mtnN* and *adeD* in the *E. coli* BL21-CodonPlus (DE3)-RIL strain (SI Table S2). The main bacterial culture for recombinant protein production was inoculated from a 30-ml overnight culture and was grown in 1-l erlenmeyer flasks containing 300 ml of 2x YT media supplemented with 100 μg ml^−1^ ampicillin and 30 μg ml^−1^ chloramphenicol at 37°C and 220 rpm for 3 h until the cell culture reached an OD_600_ of 0.6 to 0.8. The expression was then induced by adding IPTG to a final concentration of 1 mM and the cells were further grown at 37°C and 220 rpm for 3 h. For harvest, cells were pelleted by centrifugation in a Sorvall SLA-1500 fixed rotor at 4°C and 15,180xg for 10 min. The supernatant was discarded, and the pellets were flash-frozen in liquid N_2_ and stored at −80°C. On the day of protein purification, 30 ml of lysis buffer (50 mM NaH_2_PO_4_, 300 mM NaCl, 1 mg ml^−1^ lysozyme, pH 7.5) was added to the pellet while thawing on ice for approximately 30 min. Cell lysis was done by ultrasonication (Processor and Tip from Hielscher Ultrasonics, Teltow, DE) on ice 4 times for 1 min at an amplitude of 100. The supernatant was applied to a 2-ml column of Ni-NTA-agarose (Qiagen, Hilden, DE) at a flow rate of 1 ml min^−1^. The column was washed with 25 ml wash buffer (50 mM NaH_2_PO_4_ pH 7.5, 300 mM NaCl, 40 mM imidazole, pH 8) and the protein eluted in 2.5 ml elution buffer (50 mM NaH_2_PO_4_ pH 7.5, 300 mM NaCl, 250 mM imidazole, pH 8), all with a flow rate of 1 ml min^−1^. The elution was applied to a PD-10 column, which was previously equilibrated to the respective storage buffer (MetK; 100 mM Tris/HCl, 1 mM EDTA, pH 8; MtnN: 50 mM potassium phosphate, pH 7; AdeD: 50 mM Tris/HCl, 1 mM DTT, 250 mM NaCl, pH 8). The flowthrough was discarded, and the protein solution was eluted in 3.5 ml storage buffer, aliquoted and stored at −80°C.

The protocol used for production of all recombinant NAS and NAS-like proteins was conducted as described above MetK, MtnN and AdeD, with the exception that the expression was induced with 0.1 mM IPTG at 30°C for 4 h. In contrast to the purification of MetK, MtnN and AdeD, NAS and NAS-like proteins were purified via gravity flow. On the day of protein purification, 40 ml of lysis buffer (50 mM NaH_2_PO_4_, 500 mM NaCl, 50 mM Imidazole, 0.1% (v/v) Tween 20, pH 7.5, 1 tablet cOmplete™, EDTA-free Protease Inhibitor Cocktail (Roche, Mannheim, DE)) was added to the combined pellets from a total of 600 ml expression culture while thawing on ice for approximately 30 min. Cell lysis was achieved by ultrasonication (Processor and Tip from Hielscher Ultrasonics) on ice 6 times for 30 s at an amplitude of 100. The supernatant was mixed with 1 ml Ni-NTA-Agarose slurry (Qiagen) in an overhead rotator (10 rpm) at 4°C for 1 h. The suspension was applied to a column (5 ml polypropylene column, Qiagen, Hilden, Germany) with subsequent washing (5 column volumes, 50 mM NaH_2_PO_4_, 500 mM NaCl, 60 mM imidazole, 0.1 (v/v) Tween 20, pH 7.5) and elution in 2.5 ml elution buffer (50 mM NaH_2_PO_4_, 500 mM, 250 mM imidazole, pH 7.5). The eluate was applied to a PD-10 column, previously equilibrated with 1 mM Tris/HCl, pH 8, and was eluted in storage buffer (1 mM Tris/HCl, pH 8), aliquoted and stored at −80°C.

### Thin-layer chromatography (TLC) for enzyme activity testing

To assess MtnN activity, 0.1 mg ml^−1^ MtnN and 2.5 mM MTA mixed in NAS assay buffer (50 mM Tris/HCl, pH 8.7). AdeD activity was assessed by mixing 0.5 mg ml^−1^ AdeD and 3 mM adenine in NAS assay buffer. Activities of AtNAS1 mutants were assessed by mixing 0.15 mg ml^−1^ NAS, 0.1 mg ml^−1^ MtnN, 0.1 mg ml^−1^ AdeD and 5 mM SAM in NAS assay buffer. All reactions were carried out in a total volume of 30 μl at 30°C, and were stopped by flash freezing in liquid N_2_. Five μl of each reaction mixture were loaded onto a TLC plate (TLC Silica gel 60 F_254_, Merck, Darmstadt, DE). The mobile phase was 7 vol-parts of 1-propanol mixed with 8 vol-parts of H_2_O. After running for approximately 1 h in a closed TLC chamber at RT, the TLC plate was dried using a blow-dryer and documented under UV light. Additionally, the plate was sprayed with a ninhydrin solution (0.2% w/v in EtOH) using a spay bottle and incubated at 100°C for 5 min to visualize nicotianamine.

### Spectrophotometry

Spectrophotometric quantifications of NAS enzyme activities were carried out in a 96-well plate (UV-STAR® MICROPLATTE, Greiner, Frickenhausen, DE) at 37°C in a Syntex multimode reader with monitoring of light absorbance at 265 nm. Each measurement was done simultaneously in three wells (technical replicates) with a final volume of 100 μl per well. Each reaction included 1 mg ml^−1^ MtnN, 1 mg ml^−1^ AdeD, 50 mM Tris/HCl, pH 8.7, and 0.15 mg ml^−1^ NAS unless otherwise mentioned. The mixture was pre-incubated at 37°C for 3 minutes, and the enzymatic reaction started by adding 0.125 mM SAM and mixing by pipetting 10% of the total reaction volume up and down 5 times. The initial rate of change in light absorbance over time was estimated with the setting “Maximize Slope Magnitude” in ICEKAT (https://icekat.herokuapp.com/icekat), and the enzyme activity was calculated in nkat mg^−1^ protein using Lambert-Beer’s law and an extinction coefficient for adenine (λ = 265 nm) of 6,700 M^−1^ cm^−1^ (25). All NAS proteins examined in this study and their calculated molar masses can be found in SI Table S6.

### Mass spectrometry

To confirm the identity of the NAS reaction product NA in our reaction mixtures, LC-HRMS measurements were carried out using a compact™ Q-TOF (Bruker Daltonik GmbH, Life Sciences, Bremen, Germany) using the standard electrospray ionization source (positive mode). A short NUCLEODUR C18 Isis column (Macherey & Nagel, 3/2, 1.8 μm) was used for crude chromatographic separation using water/acetonitrile as mobile phase (flow rate: 0.3 ml min^−1^; eluent A: H_2_O/0.1% (v/v) HCOOH; eluent B: acetonitrile/0.1% (v/v) HCOOH; isocratic, 30% A/70% B) on an Ultimate 3000 HPLC System (consisting of a pump, autosampler, column oven and UV detector). For internal calibration we used a lock mass of *m/z* = 622.02896 (Hexakis(1H,1H,2H-perfluoroethoxy)phosphazene) and sodium formate clusters.

Reaction mixtures containing NAS were set up as described above in “TLC for enzyme activity testing”. After the reactions, one volume of 100% EtOH was added to the reaction mixture, followed by incubation at −80°C for 2 h, to precipitate any NAS in the sample. Samples were then centrifuged at 15,115×*g* and 4°C for 30 min, the supernatant transferred into a new reaction tube and dried in a rotating vacuum concentrator (Genevac Quattro miVac concentrator, Schlee GmbH, Witten, Germany). The pellet was dissolved in 120 μl ultrapure water containing 0.1% (v/v) formic acid (LC-MS grade) and stored at −20°C. Samples were injected directly into the LC-HRMS system. MS spectra were analyzed for compounds eluting at retention times between 0.31 to 0.59 min.

## Data availability

All data are contained in this manuscript and associated Supporting Information.

## Supporting Information

This article contains Supporting Information Figures S1 to S9, Tables S1 to S6 and Data S1.

## Acknowledgments

We thank Pascal Arnoux (CEA, Saint Paul lez Durance, France) for fruitful discussions and for materials. We thank Dr. Heike Holländer Czytko and Charlotte Kürten (Molecular Genetics and Physiology of Plants, Ruhr University Bochum) for preparatory work that led to this study. We thank Eckhard Hofmann (Protein Crystallography, Ruhr University Bochum) and Research Training Group 2341 “Microbial Substrate Conversion” for discussions.

## Author contributions

Conceptualization: UK, MP. Data curation: HS, MP, UK. Experimentation: HS, GR, FS. Funding acquisition: UK. Methodology: HS, MP, GR and all authors. Project administration: UK. Resources: MB, MP, UK, FS. Supervision: UK, MP. Validation: HS, GR, MP, UK, FS; Visualization: HS, MP, UK; Writing – original draft: HS, with contributions from MP, UK and FS; Writing – reviewing & editing: UK, HS, MP, FS.

## Funding and additional information

This work was funded by the German Research Foundation, Research Training Group “Microbial Substrate Conversion (MiCon)” (RTG 2341).

## Conflict of interest

The authors declare no conflict of interest.

## Notes

### Competing Interest Statement

The authors have declared no competing interest.

