## Supplemental Information pdf file for "Arabidopsis Nicotianamine Synthases (NAS) comprise a common core-NAS domain fused to a variable auto-inhibitory C-terminus"

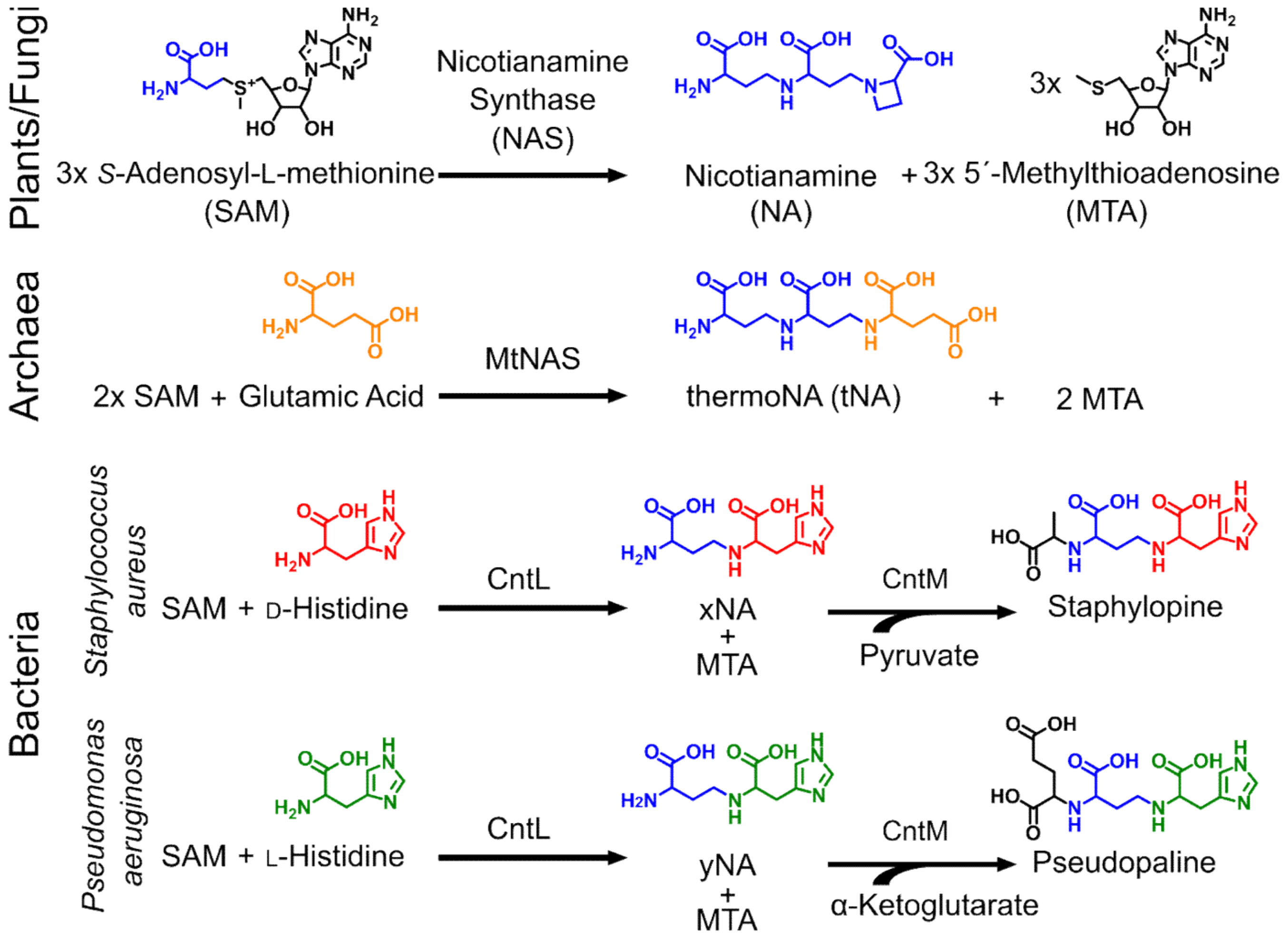


**Figure S1. Reactions catalyzed by Nicotianamine Synthase (NAS) and NAS-like proteins.**
NAS of plants and fungi utilize the α-aminobutyrate group (blue) of each of three *s*-adenosylmethionine (SAM) molecules to form one molecule of nicotianamine (NA) and three methylthioadenosine (MTA) molecules. Known NAS-like proteins of archaea appear to use one glutamate (orange) and the α-aminobutyrate groups of two SAM molecules to form one thermo-nicotianamine (tNA) and two MTA molecules. The characterized bacterial NAS-like proteins (CntL) use one D-/L-histidine (red/green) and one α-aminobutyrate group from SAM to form one xNA/yNA and MTA. Subsequently, Opine Dehydrogenase (CntM) utilizes NADPH and an α-keto acid to convert xNA with pyruvate to staphylopine, or yNA with α-ketoglutarate to pseudopaline. NAS and the NAS-like MtNAS are thought to catalyze the formation of their products from the substrates sequentially in a step-wise manner inside a central reaction cavity of the protein, from the right to the left of the products shown here.

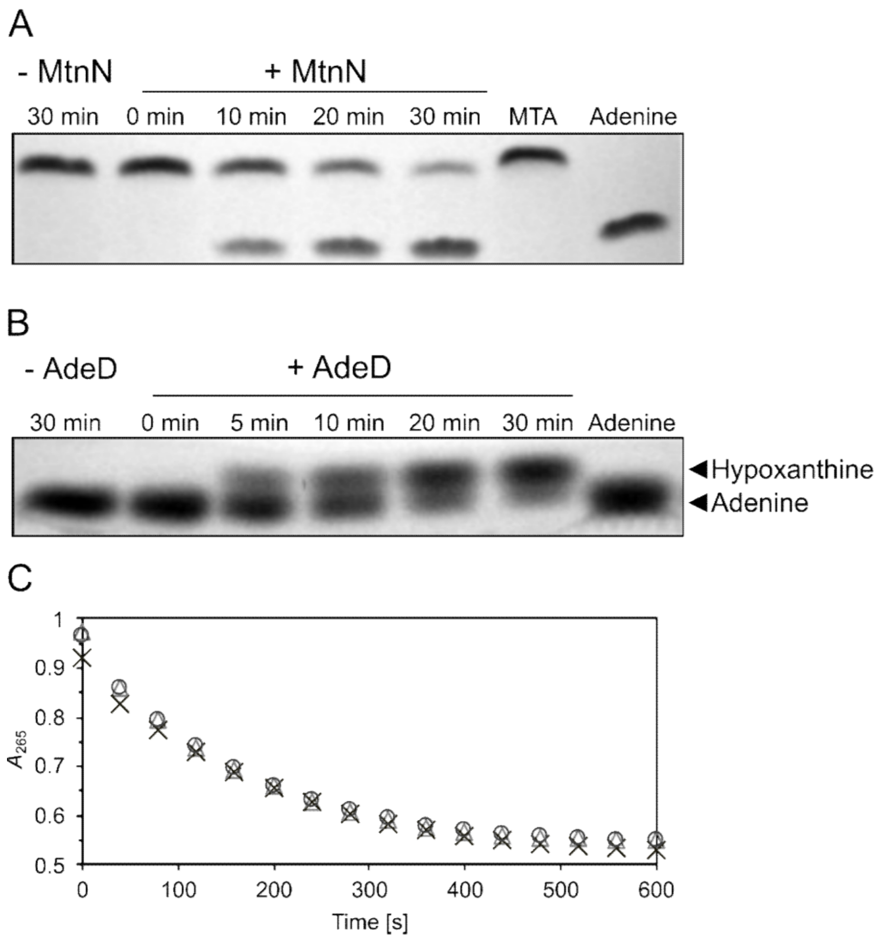


**Figure S2. Activities of purified Methylthioadenosine Nucleosidase (MtnN) and Adenine Deaminase (AdeD) in NAS reaction buffer.**
(A, B) MtnN (A) and AdeD (B) were incubated with their respective substrate in NAS reaction buffer (50 mM Tris/HCl, pH 8.7), and the reaction products were separated by thin-layer chromatography and visualized under UV light at different time points after the start of the reaction (0 min). (C) Photometric measurement over time of the coupled conversion of MTA to hypoxanthine catalyzed by the enzymes MtnN and AdeD in combination in NAS reaction buffer. Square, triangle and diamond each represent one technical replicate. MTA was added at time point 0 s.

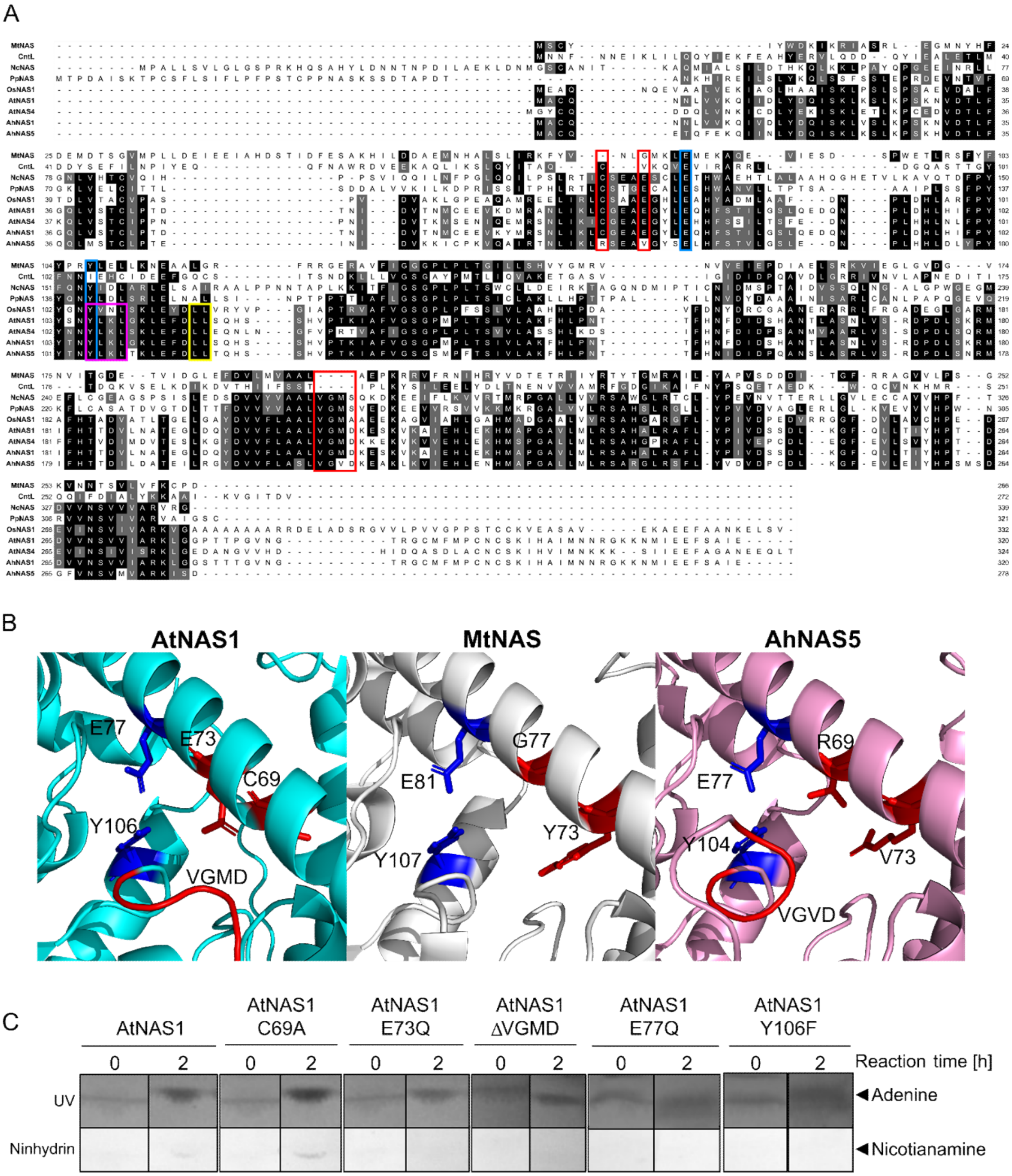


**Figure S3. Activities of purified AtNAS1 and AtNAS1 mutants.**
(A) Amino acid alignment of selected full-length nicotianamine synthases (NAS). Amino acids are shown on a black/grey background whenever ≥ 50% of them are identical/similar. Similarity groups were based on the default classification of MultipleAlignShow (ILV, FWY, KRH, DE, GAS, P, C, TNQM) (50). Blue boxes mark positions thought to be essential for the overall reaction mechanism of both NAS and NAS-like proteins, red boxes mark positions near the reaction cavity that are conserved among NAS that produce NA. A yellow and a pink box, respectively, mark an LL motif reported as essential for the *in vitro* activity of OsNAS2 and a YxxΦ motif proposed to be required for the *in planta* activity of OsNAS2 (41). (B) Protein model of AtNAS1 (generated by AlphaFold 2), MtNAS (PDB:3FPE) and AhNAS5 (generated by swissmodel) with highlighted amino acids mentioned in (A) (39). (C) AtNAS1 (4.5 µg) and MtnN (10 µg) were co-incubated in the presence of *S*-adenosyl-L-methionine (5 mM) in a total volume of 30 µl. The reaction was started by incubating the mixture at 30°C, and it was stopped immediately (0 h) or 2 h after the start of reaction by flash-freezing. Aliquots of 5 µl per reaction were separated by TLC, and the products were visualized by UV light (adenine) or ninhydrin staining (nicotianamine).

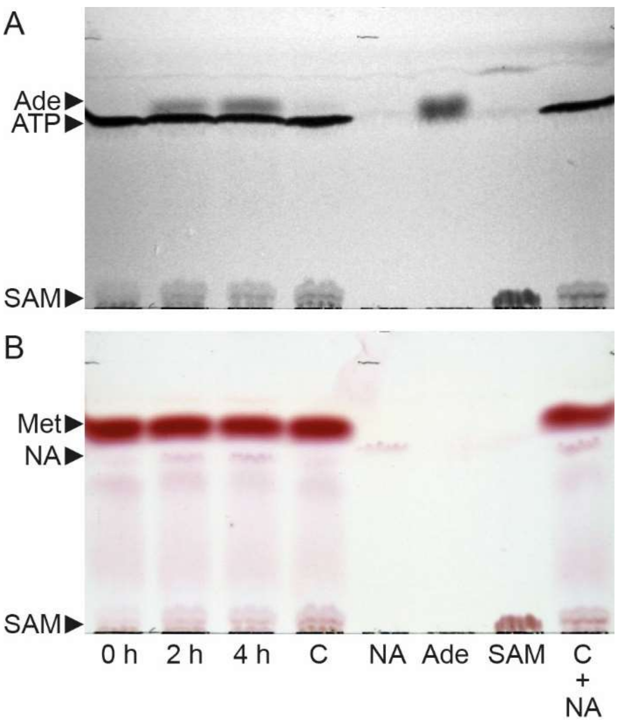


**Figure S4. One-pot biosynthesis of nicotianamine from L-methionine and ATP.**
(A, B) AtNAS1 (42 µg), MetK (60 µg) and MtnN (12 µg) where co-incubated in the presence of L-methionine (Met, 10 mM) and ATP (10 mM) in a total volume of 120 µl at 30°C for up to 4 h. At the indicated time points, 5-µL aliquots of the reaction mix were separated by TLC, and compounds were visualized by UV light (A) and ninhydrin staining (B). Adenine (Ade, 16 nmol standard), nicotianamine (NA, 8 nmol standard) and *S*-adenosylmethionine (SAM, 25 nmol standard) were run as standards. C: Negative control (conducted with heat-denatured AtNAS1), C + NA: As C, but with added nicotianamine (192 nmol).

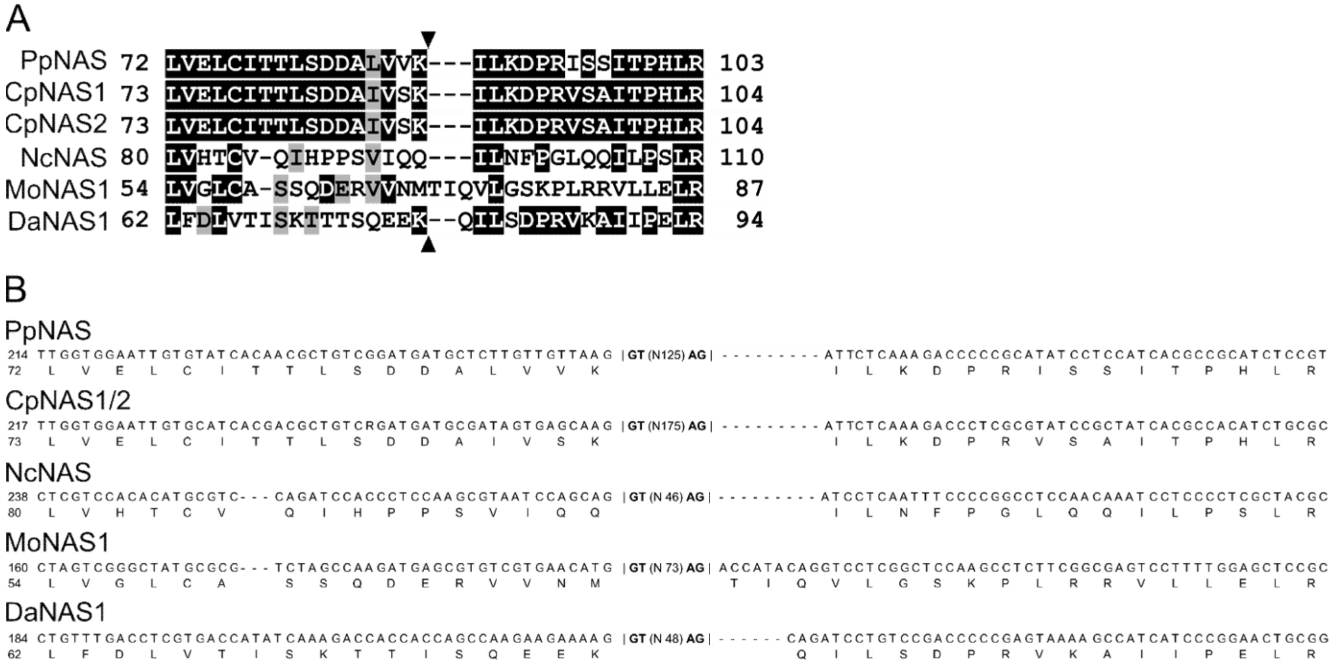


**Figure S5. *NAS* genes of the mosses *P. patens* and *C. purpureus* and of the fungi *N. crassa*, *M. oryzae* and *D. ampelina* all contain an intron at a conserved position.**
(A) To visualize the conserved position of the intron, part of an amino acid alignment is shown. The intron position is indicated by triangles. (B) Genomic sequence (upper rows) alongside amino acid sequence (bottom rows) indicated centrally within each codon of the respective *NAS* gene. The sequences are aligned as in (A). In all 6 sequences, the intron is in phase 0 (intron position given by vertical lines; N followed by a number specifies the length of each intron, and only the initial and final two nucleotides of each intron are specified). The *NAS* gene of *M. oryzae* contains a second intron downstream in its sequence. Amino acids are shown on a black/grey background whenever ≥ 50% of them are identical/similar. Similarity groups were based on the default classification of MultipleAlignShow (ILV, FWY, KRH, DE, GAS, P, C, TNQM). Pp: *Physcomitrium patens* (Pp3c8_970), CepurR40: *Ceratodon purpureus* male isolate R40 (CepurR40.10G154600), CepurGG1: *C. purpureus* female isolate GG1 (CepurGG1.10G14960), Nc.: *Neurospora crassa* (XP_958379.1), Mo: *Magnaporthe oryzae* (XP_003719353.1), Da: *Diaporthe ampelina* (KKY38707.1).

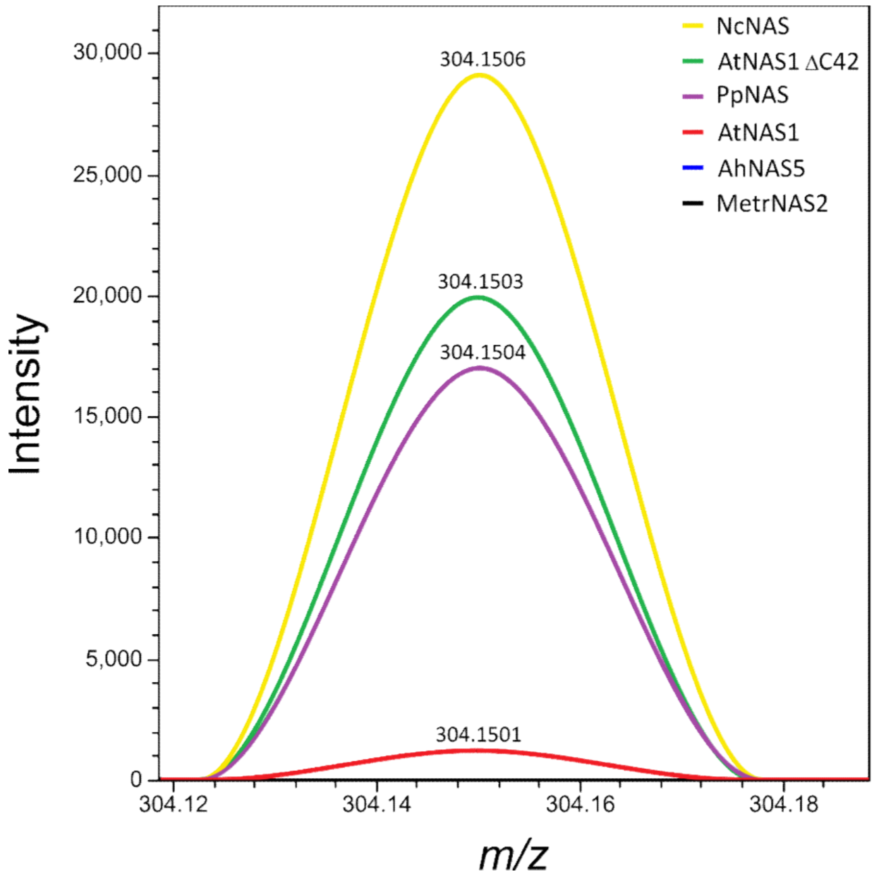


**Figure S6. Detection of nicotianamine with mass spectrometry of reaction mixtures containing different NAS.**

Detection of nicotianamine by mass spectrometry of reaction mixtures containing different NAS. Purified NAS enzymes (0.15 mg protein $ml^{-1}$) were incubated with MtnN (1 mg protein $ml^{-1}$) and 3 mM SAM in a total volume of 30 µL at 30°C for 2 h before LC-MS. HRMS was carried out in positive ion mode (for details see SI Fig. S9).

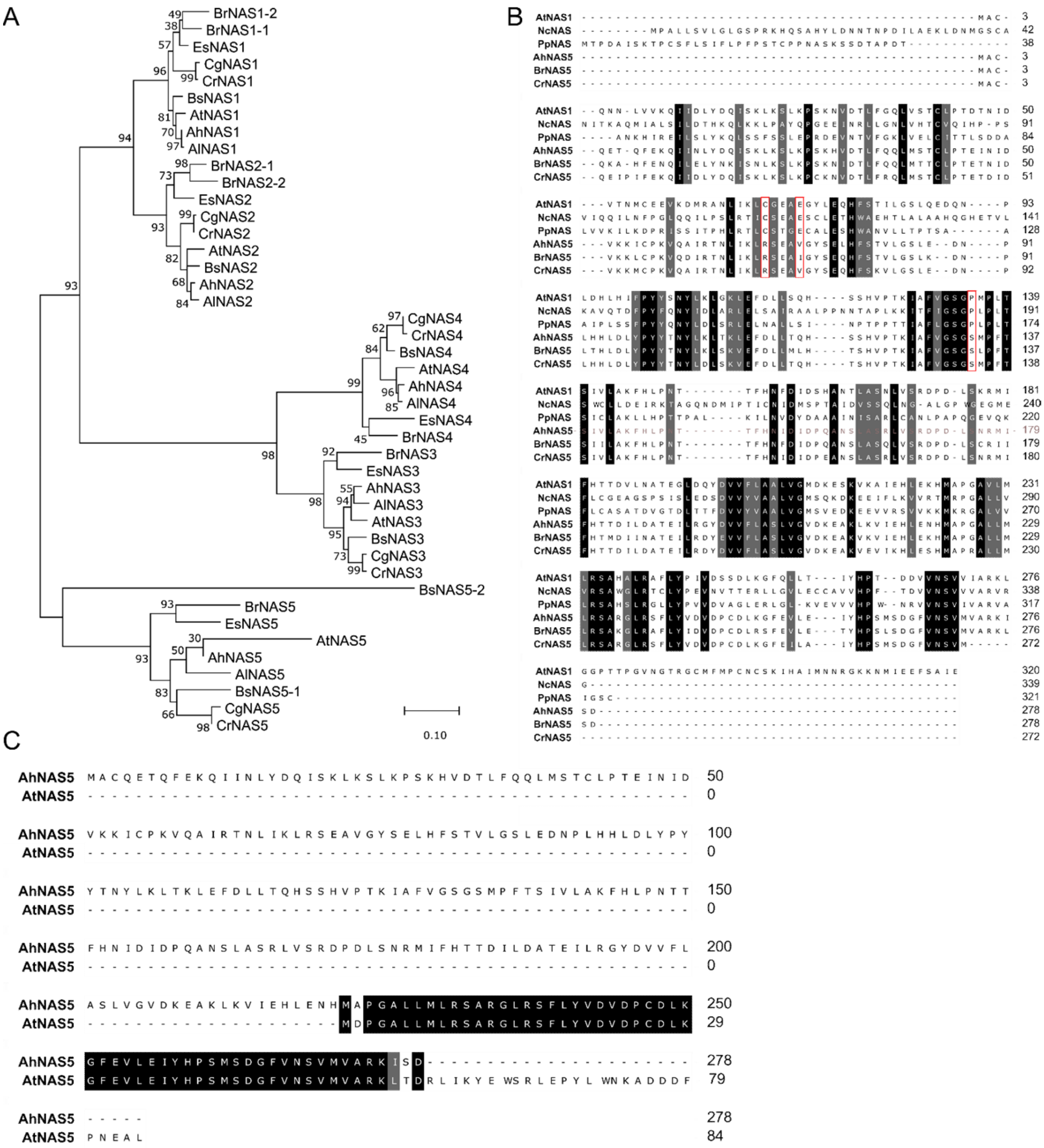


**Figure S7. NAS isoforms in the Brassicaceae family of dicotyledonous angiosperms.**
(A) Maximum likelihood phylogenetic tree of selected NAS sequences from the Brassicaceae. The existence of 5 paralogous NAS proteins in the Brassicaceae is evident. The percentage of trees in which the associated taxa clustered together is shown next to the branches. Branch lengths reflect the number of substitutions per site, scaled as indicated below the tree. (B) Full amino acid sequence alignment of selected NAS proteins observed to exhibit enzyme activity here (NcNAS, PpNAS, AtNAS1) and NAS5 proteins from *A. halleri* (no detectable enzyme activity; see SI Fig. S6), *B. napus* and *C. rubella*. Amino acids are shown on a black/grey background whenever ≥ 50% of them are identical/similar. Similarity groups were based on the default classification of MultipleAlignShow (ILV, FWY, KRH, DE, GAS, P, C, TNQM) (50). Red boxes highlight amino acids conserved among active NAS proteins which are divergent in NAS5 proteins of the Brassicaceae. (C) Full amino acid sequence alignment of NAS5 proteins from *A. halleri* and *A. thaliana. AtNAS5* (AT4G26483) is a pseudogene encoding merely a short fragment of the NAS5 protein.

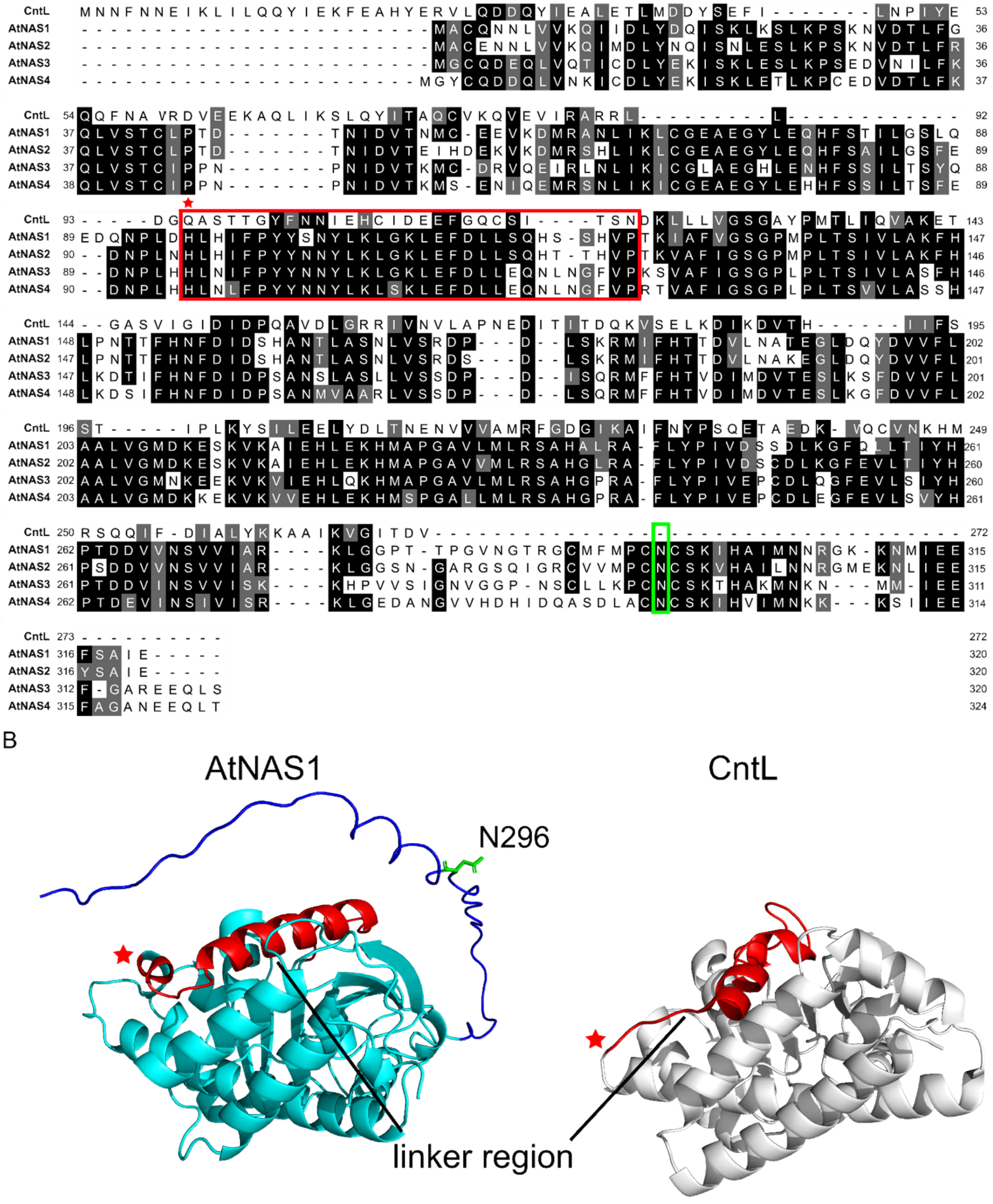


**Figure S8. Linker region in AtNAS and CntL**
(A) Full amino acid alignment of AtNAS1-4 and *Staphylococcus aureus* CntL. Red/green boxes mark the amino acids corresponding to the conformationally dynamic linker region of CntL (38)/N296 contributing to auto-inhibition of AtNAS1 (see Fig. 6). Amino acids are shown on a black/grey background whenever ≥ 50% of them are identical/similar. Similarity groups were based on the default classification of MultipleAlignShow (ILV, FWY, KRH, DE, GAS, P, C, TNQM) (50). (B) Modified protein model of AtNAS1 (left, generated by AlphaFold 2) and CntL (right, PDB:7C9M), with red/green used to identify regions as defined in (A) (39). The elongated C-terminus of AtNAS1 is colored in dark blue.

| Table S2. List of bacterial strains and plasmids. | |
|---|---|
| **Bacterial strain** | **Supplier** |
| XL1-Blue | Agilent Technologies, Santa Clara, US |
| BL21-CodonPlus (DE3)-RIL | Agilent Technologies, Santa Clara, US |
| BL21 DE3 pLysS | EMD Millipore, Burlington, US |
| **Plasmid** | **Supplier** |
| pGEM®-T Easy | Promega, Madison, US |
| pET101 Directional TOPO ™ | ThermoFisher Scientific, Waltham, US |
| pET-21b (+) | EMD Millipore, Burlington, US |
| pMA-RQ | ThermoFisher Scientific, Waltham, US |

**Table S4. PCR conditions.**

| Amplicon | Encoded Protein | PCR conditions |
|---|---|---|
| *mtnN* | MtnN | 95 °C 3 min, 30 × [95 °C 30 s, 70 °C 30 s, 72 °C 60 s], 72 °C 10 min |
| *adeD* | AdeD | 98 °C 3 min, 30 × [98 °C 10 s, 68 °C 30 s, 72 °C 60 s], 72 °C 5 min |
| *metK* | MetK | 95 °C 3 min, 30 × [95 °C 30 s, 66 °C 30 s, 72 °C 60 s], 72 °C 10 min |
| MTH675 | MtNAS | 98 °C 30 s, 30 × [98 °C 10 s, 72 °C 60 s], 72 °C 5 min |
| AT5G04950 | AtNAS1 | 94 °C 10 min, 35 × [93 °C 30 s, 52 °C 30 s, 72 °C 2 min], 72 °C 90min |
| | AtNAS1 C69A | 95 °C 2 min, 20x [98 °C 20 s, 55 °C 30 s, 72 °C 3,5 min], 72 °C 5 min |
| | AtNAS1 E73Q | 95 °C 2 min, 20x [98 °C 20 s, 68 °C 30 s, 72 °C 3,5 min], 72 °C 5 min |
| | AtNAS1 VGMD | 95 °C 2 min, 20x [98 °C 20 s, 52 °C 30 s, 72 °C 3,5 min], 72 °C 5 min |
| | AtNAS1 E77Q | 95 °C 30 s, 18x [98 °C 10 s, 55 °C 10, 72 °C 3,5 min], 72 °C 5 min |
| | AtNAS1 Y106F | 95 °C 30 s, 18x [98 °C 10 s, 55 °C 10, 72 °C 3,5 min], 72 °C 5 min |
| | AtNAS1 N296D | 95 °C 30 s, 18x [98 °C 10 s, 55 °C 10, 72 °C 3,5 min], 72 °C 5 min |
| AT5G56080 | AtNAS2 | 98 °C 30 s, 30 × [98 °C 10 s, 64 °C 15 s, 72 °C 30 s], 72 °C 5 min |
| | AtNAS2 ΔC43 | 98 °C 30s, 30 × [98 °C 10 s, 53,8 °C 10 s, 72 °C 30 s], 72 °C 10 min |
| AT1G09240 | AtNAS3 | 98 °C 30 s, 30 × [98 °C 10 s, 64 °C 15 s, 72 °C 30 s], 72 °C 5 min |
| | AtNAS3 ΔC42 | 98 °C 30s, 30 × [98 °C 10 s, 53,8 °C 10 s, 72 °C 30 s], 72 °C 10 min |
| AT1G56430 | AtNAS4 | 94 °C 10 min, 35 × [93 °C 30 s, 52 °C 30 s, 72 °C 2 min], 72 °C 90min |
| | AtNAS4 ΔC47 | 98 °C 30s, 30 × [98 °C 10 s, 53,8 °C 10 s, 72 °C 30 s], 72 °C 10 min |
| | AtNAS1 ΔC13 | 98 °C 30s, 30 × [98 °C 10 s, 70 °C 25 s, 72 °C 30 s], 72 °C 10 min |
| | AtNAS1 ΔC22 | 98 °C 30s, 30 × [98 °C 10 s, 67,2 °C 15 s, 72 °C 30 s], 72 °C 10 min |
| | AtNAS1 ΔC31 | 98 °C 30s, 30 × [98 °C 10 s, 67,2 °C 15 s, 72 °C 30 s], 72 °C 10 min |
| | AtNAS1 ΔC34 | 98 °C 30s, 30 × [98 °C 10 s, 72 °C 30 s], 72 °C 10 min |
| | AtNAS1 ΔC42 | 98 °C 30s, 30 × [98 °C 10 s, 72 °C 30 s], 72 °C 10 min |
| | AtNAS1 ΔC43 | 98 °C 30s, 30 × [98 °C 10 s, 72 °C 30 s], 72 °C 10 min |
| | AtNAS1 ΔC44 | 98 °C 30s, 30 × [98 °C 10 s, 72 °C 30 s], 72 °C 10 min |
| | AtNAS1 ΔC45 | 98 °C 30s, 30 × [98 °C 10 s, 72 °C 30 s], 72 °C 10 min |
| | AtNAS1 ΔC46 | 98 °C 30s, 30 × [98 °C 10 s, 72 °C 30 s], 72 °C 10 min |

**Table S6. Calculated molar mass for proteins used in this study.**

| Protein | molar mass [Da] |
|---|---|
| AtNAS1 | 35,547 |
| AtNAS1 ΔC42 | 30,936 |
| MtnN | 24,398 |
| AdeD | 63,739 |
| MetK | 41,952 |
| NcNAS | 36,928 |
| PpNAS | 34,505 |
| MetrNAS2 | 31,978 |
| AtNAS2 | 35,679 |
| AtNAS3 | 35,751 |
| AtNAS4 | 36,352 |
| AtNAS2 Δ43 | 31,004 |
| AtNAS3 Δ42 | 31,201 |
| AtNAS4 Δ47 | 31,203 |
| AtNAS1 ΔC13 | 34,069 |
| AtNAS1 ΔC22 | 32,991 |
| AtNAS1 ΔC31 | 31,974 |
| AtNAS1 ΔC34 | 31,659 |
| AtNAS1 ΔC43 | 30,878 |
| AtNAS1 ΔC44 | 30,821 |
| AtNAS1 ΔC45 | 30,708 |
| AtNAS1 ΔC46 | 30,580 |
| AtNAS1 N296D | 35,548 |

https://web.expasy.org/protparam/
