## Supplemental Information Figure S9 for "Arabidopsis Nicotianamine Synthases (NAS) comprise a common core-NAS domain fused to a variable auto-inhibitory C-terminus"

**Settings used in Mass Spectrometry method.**

#### Method Set: D:\Methods\Susanna\110-1300 autoMSMS pos_.m

BRUKER

### otofControl

#### General Information

| | | | |
|---|---|---|---|
| **Method Name:** | 110-1300 autoMSMS pos_.m | **Saved:** | 2018/09/05 09:38:08+02:00 |
| **Application Name:** | Bruker otofControl | **Application Version:** | 4.1.3.5 |
| **Device Type:** | compact | **Device Serial Number:** | 8255754.20147 |
| **Operator:** | Demo User | **Host:** | COMPACT-20147 |
| **Operating System:** | Windows 7 Professional | **Organisation:** | Bruker Daltonik GmbH |

#### Chromatogram

##### Chromatogram Traces

| Enabled | Color | Type | Masses | Width | Polarity | Filter |
|---|---|---|---|---|---|---|
| On | Red | BPC | | | ± | MS |
| On | Blue | TIC | | | ± | MS |
| On | Black | TIC | | | ± | All MS/MS |

#### SPL

| | |
|---|---|
| **Scheduled Precursor List:** | Off |

#### Segment 1

#### 0 .... 0.02 min

##### Main

| | | | |
|---|---|---|---|
| **Polarity:** | Positive | **Scan Mode:** | MS |
| **Mass Range from:** | 110 m/z | **Mass Range to:** | 1300 m/z |
| **Rolling Average:** | Off | **Rolling Average No.:** | 2 |
| **Spectra rate:** | 8.00 Hz | **View:** | Expert |

##### Mode

| | | | |
|---|---|---|---|
| **Save Spectra:** | Line and Profile Spectra | **Line Spectra Calculation:** | Use Maximum Intensity |
| **Absolute Threshold (per 1000 sum.):** | 25 cts. | **Peak Summation Width:** | 3 pts. |
| | | **Focus Active:** | Off |
| **Mark as Calibration Segment:** | Off | | |

##### Source

| | | | |
|---|---|---|---|
| **Source:** | ESI | | |
| **End Plate Offset:** | 500 V | **Capillary:** | 4500 V |
| **Nebulizer:** | 2.2 Bar | **Dry Gas:** | 10.0 l/min |
| **Dry Temp:** | 220 °C | **Divert Valve:** | Waste 1-6 |

##### Tune

| | | | |
|---|---|---|---|
| **Funnel 1 RF:** | 150.0 Vpp | **Funnel 2 RF:** | 200.0 Vpp |
| **isCID Energy:** | 0.0 eV | **Hexapole RF:** | 50.0 Vpp |
| **Ion Energy:** | 4.0 eV | **Low Mass:** | 90.0 m/z |
| **Collision Energy:** | 7.0 eV | **Pre Pulse Storage:** | 5.0 µs |
| **Stepping:** | On | **Mode:** | Basic |
| **Collision RF from:** | 550.0 Vpp | **Collision RF to:** | 550.0 Vpp |
| **Transfer Time from:** | 80.0 µs | **Transfer Time to:** | 80.0 µs |
| **Timing from:** | 50 % | **Timing to:** | 50 % |
| **Collision Energy from:** | 100 % | **Collision Energy to:** | 250 % |
| **Timing from:** | 50 % | **Timing to:** | 50 % |

### MS/MS

| | |
|---|---|
| **Auto MS/MS:** | Off |

##### *MRM*

| | |
|---|---|
| **MRM:** | Off |

##### *isCID*

| | |
|---|---|
| **isCID (MS-MS/MS):** | Off |

##### *bbCID*

| | |
|---|---|
| **bbCID (MS-MS/MS):** | Off |

#### Segment 2
#### 0.02 .... 0.3 min

##### Main

| | | | |
|---|---|---|---|
| **Polarity:** | Positive | **Scan Mode:** | MS |
| **Mass Range from:** | 110 m/z | **Mass Range to:** | 1300 m/z |
| **Rolling Average:** | Off | **Rolling Average No.:** | 2 |
| **Spectra rate:** | 8.00 Hz | **View:** | Expert |

##### Mode

| | | | |
|---|---|---|---|
| **Save Spectra:** | Line and Profile Spectra | **Line Spectra Calculation:** | Use Maximum Intensity |
| **Absolute Threshold (per 1000 sum.):** | 25 cts. | **Peak Summation Width:** | 3 pts. |
| | | **Focus Active:** | Off |
| **Mark as Calibration Segment:** | On | | |

##### Source

| | | | |
|---|---|---|---|
| **Source:** | ESI | | |
| **End Plate Offset:** | 500 V | **Capillary:** | 4500 V |
| **Nebulizer:** | 2.2 Bar | **Dry Gas:** | 10.0 l/min |
| **Dry Temp:** | 220 °C | **Divert Valve:** | Source 1-2 |

##### Tune

| | | | |
|---|---|---|---|
| **Funnel 1 RF:** | 150.0 Vpp | **Funnel 2 RF:** | 200.0 Vpp |
| **isCID Energy:** | 0.0 eV | **Hexapole RF:** | 50.0 Vpp |
| **Ion Energy:** | 4.0 eV | **Low Mass:** | 90.0 m/z |
| **Collision Energy:** | 7.0 eV | **Pre Pulse Storage:** | 5.0 µs |
| **Stepping:** | On | **Mode:** | Basic |
| **Collision RF from:** | 550.0 Vpp | **Collision RF to:** | 550.0 Vpp |
| **Transfer Time from:** | 80.0 µs | **Transfer Time to:** | 80.0 µs |
| **Timing from:** | 50 % | **Timing to:** | 50 % |
| **Collision Energy from:** | 100 % | **Collision Energy to:** | 250 % |
| **Timing from:** | 50 % | **Timing to:** | 50 % |

### MS/MS

| | |
|---|---|
| **Auto MS/MS:** | Off |

##### *MRM*

| | |
|---|---|
| **MRM:** | Off |

##### *isCID*

| | |
|---|---|
| **isCID (MS-MS/MS):** | Off |

##### *bbCID*

| | |
|---|---|
| **bbCID (MS-MS/MS):** | Off |

#### Segment 3

#### 0.3 .... unlimited min

##### Main

| | | | |
|---|---|---|---|
| **Polarity:** | Positive | **Scan Mode:** | Auto MS/MS |
| **Mass Range from:** | 110 m/z | **Mass Range to:** | 1300 m/z |
| **Rolling Average:** | Off | **Rolling Average No.:** | 2 |
| **Spectra rate:** | 8.00 Hz | **View:** | Expert |

##### Mode

| | | | |
|---|---|---|---|
| **Save Spectra:** | Line and Profile Spectra | **Line Spectra Calculation:** | Use Maximum Intensity |
| **Absolute Threshold (per 1000 sum.):** | 25 cts. | **Peak Summation Width:** | 3 pts. |
| | | **Focus Active:** | Off |
| **Mark as Calibration Segment:** | Off | | |

##### Source

| | | | |
|---|---|---|---|
| **Source:** | ESI | | |
| **End Plate Offset:** | 500 V | **Capillary:** | 4500 V |
| **Nebulizer:** | 2.2 Bar | **Dry Gas:** | 10.0 l/min |
| **Dry Temp:** | 220 °C | **Divert Valve:** | Waste 1-6 |

#### Tune

| | | | |
|---|---|---|---|
| **Funnel 1 RF:** | 150.0 Vpp | **Funnel 2 RF:** | 200.0 Vpp |
| **isCID Energy:** | 0.0 eV | **Hexapole RF:** | 50.0 Vpp |
| **Ion Energy:** | 4.0 eV | **Low Mass:** | 90.0 m/z |
| **Collision Energy:** | 7.0 eV | **Pre Pulse Storage:** | 5.0 µs |
| **Stepping:** | On | **Mode:** | Basic |
| **Collision RF from:** | 550.0 Vpp | **Collision RF to:** | 550.0 Vpp |
| **Transfer Time from:** | 80.0 µs | **Transfer Time to:** | 80.0 µs |
| **Timing from:** | 50 % | **Timing to:** | 50 % |
| **Collision Energy from:** | 100 % | **Collision Energy to:** | 250 % |
| **Timing from:** | 50 % | **Timing to:** | 50 % |

## MS/MS

| | | | |
|---|---|---|---|
| **Auto MS/MS:** | On | | |
| **Precursor Ion List:** | Exclude | **Cycle Time:** | 0.5 sec |
| **Threshold (per 1000 sum.) Absolute:** | 400 cts | **Active Exclusion:** | On |
| | | **Exclude after:** | 3 Spectra |
| | | **Reconsider Precursor:** | On |
| **Release after:** | 0.20 min. | **Smart Exclusion:** | Off |
| **if Curent Intens./Prev. Intens.:** | 1.8 | | |
| **Smart Exclusion:** | 2 x | | |

##### Exclude Mass List

| Mass Range Start | Mass Range End | |
|---|---|---|
| 102.08 | 102.18 | 1 |
| 621.98 | 622.08 | 2 |
| 643.96 | 644.06 | 3 |
| 659.94 | 660.04 | 4 |

#### *Auto MS/MS Preference*

| | | | |
|---|---|---|---|
| **Preferred Range:** | Off | | |
| **Preferred Range Low:** | 2 | **Preferred Range High:** | 5 |
| **Exclude Singly:** | Off | **Exclude unknown:** | Off |
| **Group Length:** | 3 | **Strict Active Exclusion:** | Off |
| **Sort Precursors by:** | Intensity | **Preferred mass list:** | Empty list |

#### *Auto MS/MS Multi CE*

**Auto MS/MS Multi CE:** Off

#### *SILE*

**SILE:** Off

#### *CID*

**Fallback Charge State:** 1 z

##### *Isolation + Fragmentation List*

| Type | Mass [m/z] | Width [m/z] | Collision Energy [eV] | Charge State | |
|---|---|---|---|---|---|
| Base | 100.00 | 4.00 | 20.0 | 1 | 1 |
| Base | 500.00 | 5.00 | 20.0 | 1 | 2 |
| Base | 1000.00 | 6.00 | 20.0 | 1 | 3 |
| Base | 1300.00 | 8.00 | 30.0 | 1 | 4 |

#### *CID Acquisition*

| | | | |
|---|---|---|---|
| **Acquisition:** | On | **Spectra Rate MS:** | 8.00 Hz |
| **MS/MS low (per 1000 sum.):** | 10000.0 cts. | **MS/MS low:** | 1600 x |
| **MS/MS high:** | 100000.0 cts. | **MS/MS high:** | 800 x |
| **Total Cycle Time Range:** | n/a sec | **Absolute Threshold :** | n/a cts. |

#### *MRM*

**MRM:** Off

#### *isCID*

**isCID (MS-MS/MS):** Off

#### *bbCID*

**bbCID (MS-MS/MS):** Off
